# DPM1 through SERPINB5 modulates desmosomal adhesion and epidermal differentiation

**DOI:** 10.1101/2022.12.28.522133

**Authors:** Maitreyi Rathod, Henriette Franz, Vivien Beyersdorfer, Marie-Therès Wanuske, Karen Leal Fischer, Pauline Hanns, Chiara Stüdle, Aude Zimmermann, Katarzyna Buczak, Camilla Schinner, Volker Spindler

## Abstract

Glycosylation is essential to facilitate cell-cell adhesion and differentiation. We determined the role of the dolichol phosphate mannosyltransferase (DPM) complex, a central regulator for glycosylation, for desmosomal adhesive function and epidermal differentiation. Deletion of the key molecule of the DPM complex, DPM1, in human keratinocytes resulted in weakened cell-cell adhesion, impaired localization of the desmosomal components desmoplakin and desmoglein-2, and led to cytoskeletal organization defects in human keratinocytes. In a 3D organotypic human epidermis model, loss of DPM1 caused impaired differentiation with abnormally increased cornification, reduced thickness of non-corneal layers, and formation of intercellular gaps in the epidermis. Using proteomic approaches, SERPINB5 was identified as DPM1-dependent interaction partner of desmoplakin. Mechanistically, SERPINB5 reduced desmoplakin phosphorylation at serine 176, which was required for strong intercellular adhesion. These results uncover a novel role of the DPM complex in linking desmosomal adhesion with epidermal differentiation.

## INTRODUCTION

Desmosomes are vital mediators of intercellular adhesion and are mechanistically linked to several diseases, such as the blistering skin disease pemphigus, arrhythmogenic cardiomyopathy and different cancer entities^1–3^. Desmosomes are highly ordered complexes comprising of adhesion molecules and plaque proteins. Desmogleins (DSG) 1-4 and desmocollins (DSC) 1-3 belong to the cadherin family of adhesion molecules, which interact in a calcium-dependent manner^4,5^. While desmosomal cadherins mediate cell-cell contact by means of their extracellular domains, the intracellular domains interact with the armadillo proteins plakophilin 1-3 and plakoglobin. These connect to desmoplakin (DSP), which links the entire complex to intermediate filaments of the cytoskeleton, thereby providing mechanical strength to cells and tissues^6^. In addition to intermediate filaments, the actin cytoskeleton also has implications on desmosome assembly, where actomyosin contractility and the cortical actin network are required for membrane translocation and mobility of desmosomal proteins^7^. Cell-cell adhesion not only provides mechanical strength and tissue integrity, but also influences differentiation processes. A well-coordinated balance of cell-cell adhesion via desmosomes, adherens junctions and tight junctions, as well as differentiation of keratinocytes into the cornified layer is important to maintain barrier functions of the epidermis^8^. Alterations in several desmosomal components have led to barrier and differentiation defects of epidermis, highlighting the importance of desmosomal cadherins for maintaining tissue integrity and mediating a proper differentiation process^9–13^. However, the precise underlying mechanisms are only partially understood.

Various post translational modifications (PTMs) seem to play an important role in mediating the subcellular localization of desmosomal proteins, and glycosylation serves as an important PTM in this context. Studies have shown the impact of glycosylation in maintaining tissue homeostasis, protein trafficking, cell signaling, proliferation, differentiation, and cell adhesion^14,15^. Glycosylation is a complex process of glycan processing and maturation with different end products (mannose rich, complex and hybrid). All glycans are built of basic units comprising mannose, N-acetylglucosamine, galactose, fucose and sialic acid^16^. The dolichol phosphate mannosyltransferase (DPM) complex is a crucial mediator in this process as it is responsible for donating mannose residues that feed into all glycosylation pathways^17–19^. Defects in DPM1 lead to congenital glycosylation disorders and seizures^20^. With regard to desmosome function, it has been shown that inhibiting N-linked glycosylation results in impaired desmosome formation and stability^21^. It has also been demonstrated that N-linked glycosylation at multiple sites is responsible for the incorporation of DSC2 in the plasma membrane^22^. Further, O-linked glycosylation of PG at the N-terminus protects it from proteolytic degradation and enhances cell-cell adhesion^23^. Apart from its role in mediating cell-cell adhesion, glycosylation has also been studied in context of epidermal differentiation. CRISPR/Cas9-based modifications of several glycosylation genes in organotypic models of epidermis led to differentiation defects^24^. Further, glycosylation was implicated in the process of desquamation and shedding of corneocytes from the surface^25^. However, the molecular mechanisms on how glycosylation regulates cell-cell adhesion and subsequent effects on epidermal differentiation are not well understood.

Here we study the contribution of the DPM complex for desmosomal adhesion and epidermal differentiation using keratinocytes and organotypic models of the human epidermis.

## RESULTS

### DPM1 modulates cell-cell adhesion and differentiation in a human organotypic epidermis model

Glycosylation is well established as an important PTM for modulating cell-cell adhesion. We first aimed to understand the role of the DPM group of glycosylation modulators for regulation of desmosomal adhesion. We generated CRISPR/Cas9-mediated knockout (KO) cell lines of the three DPM isoforms with respective non-targeting control lines (NT) in HaCaT keratinocytes. In line with the occurrence as a complex, loss of DPM2 or DPM3 led to depletion of DPM1 (**Fig S1a**). Dispase-based intercellular adhesion assays showed that loss of DPM1, and to milder extent loss of DPM2 and DPM3, impaired intercellular adhesion, indicated by an increased number of fragments after applying shear stress to a detached monolayer (**Fig 1a, b**). A KO cell line for the desmosomal protein DSG2 was used as a control for reduced cell-cell adhesion. CRISPR/Cas9-mediated KO of DPM1 in primary human keratinocytes also resulted in loss of cell-cell adhesion, supporting the relevance of DPM1 for keratinocyte intercellular adhesion (**Fig 1c, d**). Interestingly, biochemical characterization of desmosomal proteins in HaCaT keratinocytes showed no major changes in total protein content upon loss of the DPM complex (**Fig S1b, c**). Because DPM1 is the enzymatically active protein of the complex, we focused on DPM1 for further experiments. The human epidermis is a stratified epithelium that differentiates into several layers during its course of maturation to provide a fully functional epidermis (**Fig 1e**). Immunostaining in human foreskin tissue showed expression of DPM1 in all layers of the differentiated epidermis, suggesting a contribution of this protein to epidermal homeostasis (**Fig S1d**). We generated an organotypic 3D-reconstructed human epidermis models (3D-RHE) which reproduced the differentiation process of interfollicular epidermis as indicated by CK10 and filaggrin staining and which showed expression of DPM1 throughout all non-corneal layers (**Fig 1f, S1d, e**). Depletion of DPM1 in 3D-RHEs caused defects in the differentiation process, indicated by thickening of the corneal layer and reduced thickness of the other layers, with the total epidermal thickness being unchanged (**Fig 1f-i**). These changes were unrelated to proliferation in the basal layer under DPM1 loss, as indicated by the proliferation marker Ki67 being similar in both sgNT1 and sgDPM1 conditions (**Fig 1j**). Further, DPM1 loss was accompanied by the presence of intercellular spaces within the viable layers of the epidermis, indicating perturbed cell-cell adhesion (**Fig 1k**).

**Figure 1:**
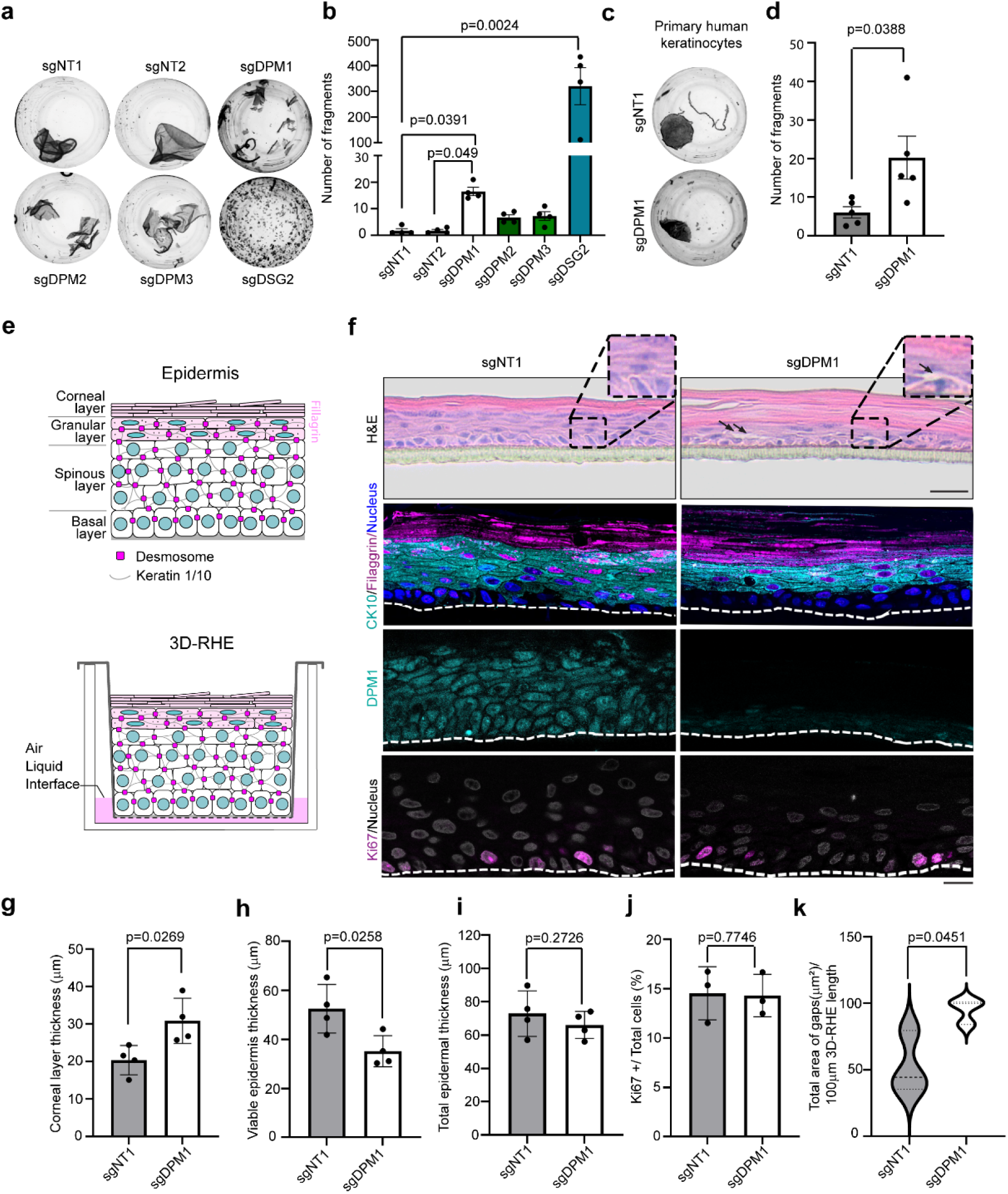
**a-b)** Dispase-based dissociation assays to semi quantitatively assess cell-cell adhesion in HaCaT keratinocytes (N=4). One-way ANOVA, Dunn’s multiple comparison test. Higher fragment numbers represent lower adhesive strength. **c-d)** Dispase-based dissociation assays of primary human keratinocytes (N=5). Unpaired Student’s t-test. **e)** Schematic of epidermis (top) and of 3D-RHE model design from primary human keratinocytes (bottom). **f)** H&E staining and immunostainings for CK10/Filaggrin, DPM1, and Ki67 of control (sgNT1) and DPM1 knockdown (sgDPM1) 3D-RHEs 12 days post airlift. Insets and arrows denote interfollicular gaps in different layers of epidermal equivalents of sgDPM1. White dashed line indicates insert membrane. Scale bar: 50µm distance on H&E images and 10µm on immunostaining images. **g-i)** Quantification of epidermal thickness parameters from 4 independent biological replicates, from 2 different donors. Each dot represents one biological replicate. Unpaired Student’s t-test. **j)** Analysis showing Ki67-positive nuclei normalised to total number of nuclei (N=3). Unpaired Student’s t-test. **k)** Violin plot showing quantification of intercellular spaces within the epidermis in sgNT1 and sgDPM1 3D-RHE. The cut off for defining intercellular spaces was set to be greater than 50µm^2^ to exclude shrinking artefacts. A minimum of 20 individual fields of view were used for analysis from 3 independent biological replicates. Unpaired Student’s t-test.

### Loss of DPM1 impairs desmoplakin surface localization and cytoskeletal arrangements in human keratinocytes

Based on the biochemical analysis, changes in the protein expression of desmosomal components and E-cadherin did not account for the reduced intercellular adhesion upon loss of DPM1. We thus investigated the localization of the essential desmosomal protein desmoplakin (DSP) in HaCaT and primary human keratinocytes. The number of DSP puncta were significantly reduced at cell borders upon loss of DPM1, indicating reduced number of desmosomes (**Fig 2a, b**). DSG2, which is a desmosomal cadherin specifically clustering in desmosomes^26^, was also reduced at the cell membrane under sgDPM1 conditions in HaCaT keratinocytes. In addition, DPM1 KO in primary human keratinocytes, reproduced the findings in HaCaT keratinocytes (**Fig 2c**). Interestingly, DSP staining appeared fragmented and more localized to the cytosol in sgDPM1 3D-RHEs compared to sgNT1 control (**Fig 2d**). With DSP serving as anchor for keratin intermediate filaments, we also tested whether the reduced DSP membrane localization was associated with changes in keratin distribution in DPM1 KO cells. Pan-cytokeratin staining showed a redistribution away from cell-cell contact sites, creating a less dense keratin network at the cell cortex. Further, individual keratin filaments appeared thickened under loss of DPM1 (**Fig 2e-g**). Depletion of DPM1 further led to redistribution of the actin cytoskeleton, with the cortical actin belt appearing more diffusely localized throughout the cytosol (**Fig 2h-j**). Changes in the cytoskeletal organisations can affect the mechanical properties of the cell^27^. Thus, we used an atomic force microscopy (AFM)-based approach to quantify the elasticity of both control and DPM1 KO HaCaT keratinocytes. DPM1 KO cells showed significantly reduced cellular elasticity, quantified as Young’s modulus (**Fig 2k**). Further, to address whether the reduced expression of DSP on the surface in DPM1 KO conditions correlates with changes in the membrane stability of DSP, we performed Fluorescence Recovery After Photobleaching (FRAP) experiments to assess the dynamics of DSP at cell-cell interfaces. FRAP showed a significant increase in the mobile fraction of DSP, indicating reduced DSP stability at cell junctions upon loss of DPM1 (**Fig 2l, m**). Together these results suggest a modulation of DSP localization and function by DPM1.

**Figure 2:**
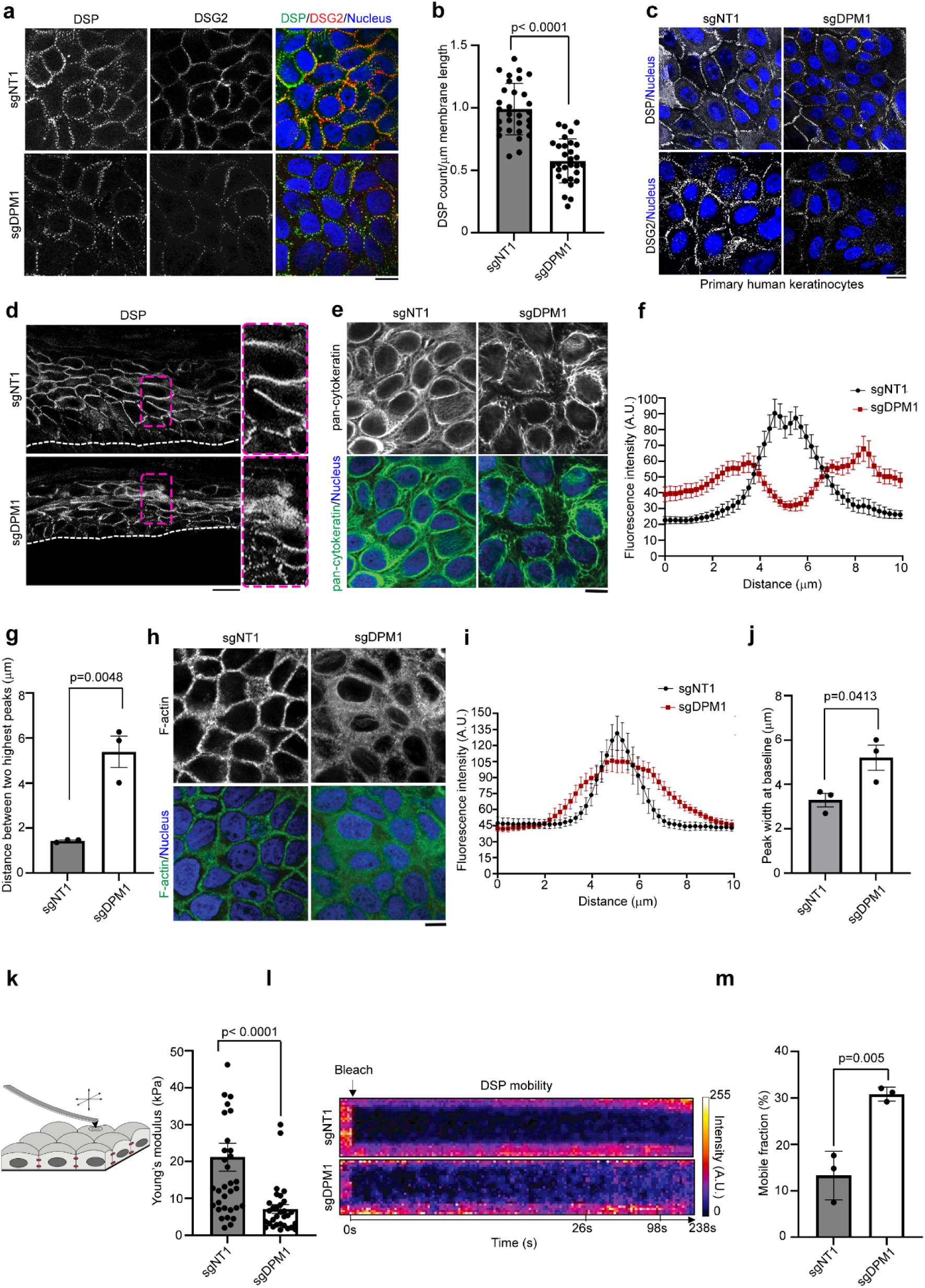
**a)** Immunostaining of desmoplakin (DSP) and desmoglein 2 (DSG2) in HaCaT keratinocytes. Merge indicates overlap between DSP, DSG2 and nuclei stained with DAPI. Scale bar:10µm distance. **b)** Quantification of the number of DSP puncta over the respective length of cell membrane (µm) from individual cells (represented by individual dots), N=3. Unpaired Student’s t-test. **c)** Immunostaining of DSP and DSG2 in primary human keratinocytes. Scale bar: 10µm distance. Panel shows representatives of 3 biological replicates. **d)** Immunostaining of DSP in 3D-RHE of control (sgNT1) and sgDPM1 conditions. White dashed line indicates insert membrane. Magenta dashed rectangles mark regions magnified on the right. Representative of 3 biological replicates. Scale bar indicates 10 µm distance. **e-g)** Keratin staining depicted by pan-cytokeratin in control and DPM1 KO HaCaT keratinocytes. Analysis done by quantifying the keratin staining intensity (A.U.) across cell borders spanning a distance of 10µm. Peak width was calculated between the 2 highest points from the distribution plot profile graph. Scale bar indicates 10 µm distance. 30 individual cells were quantified from 3 independent biological replicates. Unpaired Student’s t-test. **h-j)** F-actin stained by phalloidin in control and DPM1 KO HaCaT keratinocytes. Analysis done by quantifying the F-actin staining intensity (A.U.) across cell borders spanning a distance of 10µm. The width of the peaks was calculated between the baseline values from distribution plot profile graphs. Scale bar: 10 µm distance. 30 individual cells were analysed from 3 independent biological replicates. Unpaired Student’s t-test. **k)** Schematic of the AFM setup to measure cellular elasticity. Graph shows cellular elasticity, indicated by Young’s modulus (kPa). Each dot represents single cells from 3 biological replicates. Unpaired Student’s t-test. **l-m)** Kymographs and mobile fraction analysis derived from fluorescence-based recovery after photobleaching (FRAP) assays used to measure DSP stability at cell-cell contact sites (N=3). Each dot represents one biological replicate. Unpaired Student’s t-test.

### Proteome analysis suggest SERPINs as mediators of the changes in adhesion and differentiation upon loss of DPM1

As the DPM complex mediates essential steps of glycosylation, we tested whether loss of DPM1 results in altered glycosylation patterns of DSG2 or DSP. PNGase F-based mobility shift assay showed that DSG2 migrates at a lower molecular weight upon treatment with PNGase F (which cleaves all N-linked oligosaccharides) indicating that DSG2 is glycosylated. However, we did not observe differences in the migration pattern between sgNT1 and sgDPM1 cells. DSP showed no mobility shift upon PNGase F treatment (**Fig S2a**). This indicates that the altered localization of these molecules is independent of N-linked glycosylation. To gain more mechanistic insights into the regulatory roles of DPM1 on desmosomes, we used a global proteomics approach to identify differentially regulated proteins in DPM1 KO HaCaT keratinocytes. Here, various sets of proteins exhibited differential expression, indicating that loss of DPM1 has a profound impact on the proteome of HaCaT keratinocytes (**Fig S2b, S2c**). Glycoproteins, proteases and several proteins of the SERPIN group were significantly downregulated in DPM1 KO conditions, while various keratins, RAB GTPase and some Sec complex proteins were upregulated (**Fig 3a**). STRING-based pathway analysis of differentially expressed proteins showed significant upregulation of pathways correlating with cornification, actin filament assembly, cytoskeletal reorganization and protein folding upon loss of DPM1. Specifically, the changes in cornification and cytoskeletal organization pathways are in agreement with the structural alterations observed upon loss of DPM1. Vice versa, many metabolic pathways such as nitrogen metabolic process, carbohydrate derived metabolic process, glycosyl compound metabolic pathway, catabolic processes and purine ribonucleoside were downregulated upon loss of DPM1 (**Fig 3b**). In addition to the global changes observed, we focused on the interacting partners of DSP, since we saw major impairments in DSP surface localization and stability. DSP immunoprecipitation-based (**Fig S2d**) proteomic analysis outlined binding partners of DSP in sgNT1 or sgDPM1 cells (**Fig 3c**). Interestingly, SERPINB5 appeared as the most differential hit, with an interaction with DSP being present in control cells only. Overexpression of SERPINB5-GFP in sgNT1 cells and GFP-based pull down of the fusion protein confirmed interaction with DSP (**Fig 3d**). In line with these observations, we detected a reduced expression of SERPINB5 in DPM1 KO HaCaT keratinocytes by western blot and immunofluorescence-based assays (**Fig S2e-h**). Because global proteome data also showed several proteins of the SERPIN group downregulated in DPM1 KO cells, we focused on understanding the role of SERPINs in modulating DSP localization and cell-cell adhesion. In addition to SERPINB5, we expressed SERPINB3 and SERPINB4 as GFP fusion constructs in DPM1 KO HaCaT keratinocytes (**Fig S2i**). In adhesion assays, both SERPINB3 and SERPINB5 significantly rescued the loss of cell-cell adhesion in response to DPM1 loss, while SERPINB4 showed no protective effect (**Fig 3e, f**). In line with these results, we confirmed that expression of SERPINB5 also rescued loss of cell cohesion induced by sgDPM1 in primary human keratinocytes (**Fig S2j**).

**Figure 3:**
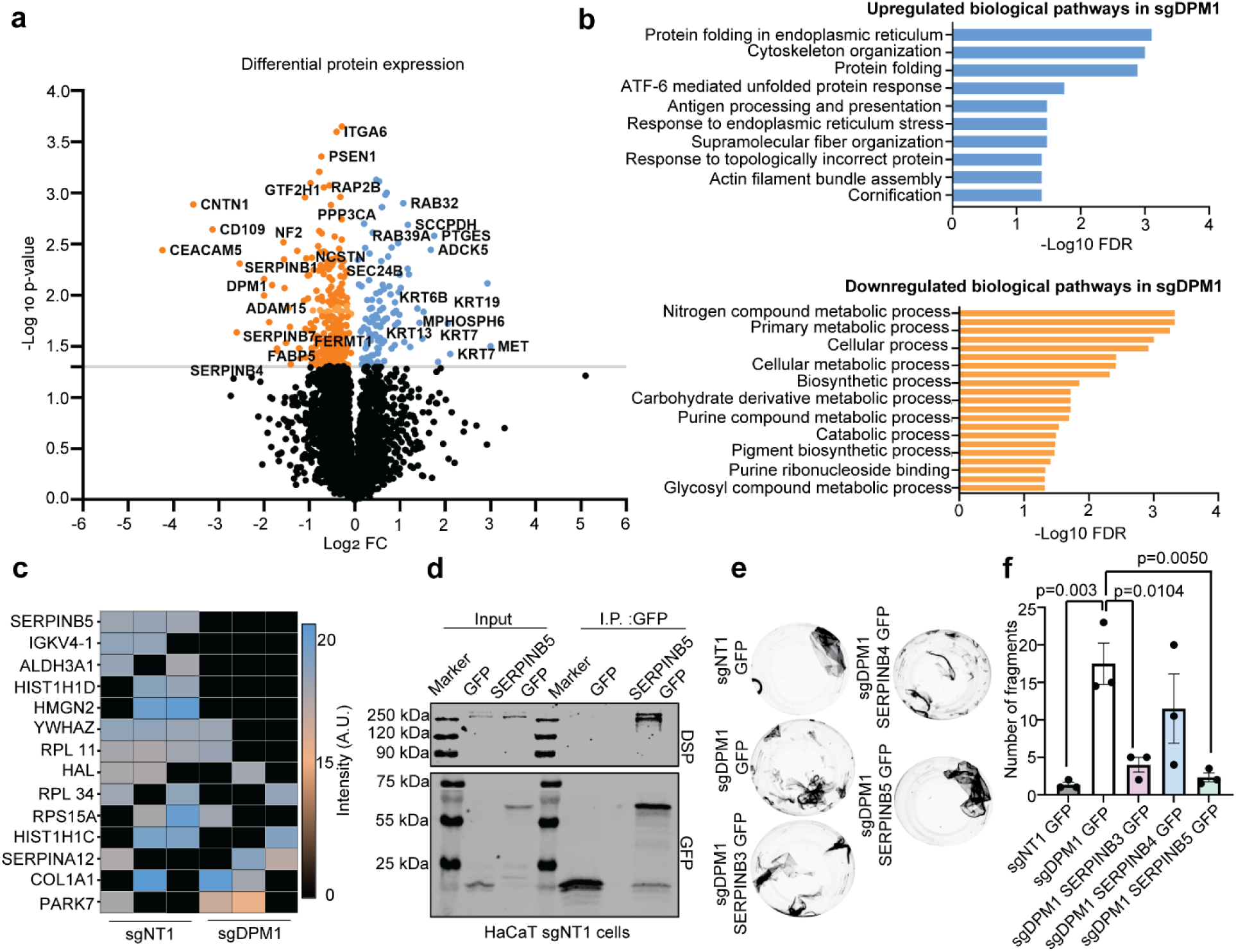
**a)** Volcano plot showing differential protein expression in sgDPM1 with respect to control sgNT1 HaCaT keratinocytes from 3 biological replicates. −Log 10 p values above 1.3 (0.05) were considered significant (marked by grey line on plot). Orange dots indicate proteins which were significantly downregulated in DPM1 KO cells, whereas blue dots indicate proteins that were significantly upregulated in DPM1 KO cells. **b)** STRING-based biological pathway analysis of significantly modulated proteins (orange: downregulated pathways and blue: upregulated pathways). X-axis denotes −log10 of false discovery rate values (FDR), with a cut off set to 0.05 for significance. **c)** Heat map showing binding partners of DSP which were absent in either sgNT1 or sgDPM1 cells in at least two biological replicates (N=3). **d)** Co-immunoprecipitation assay showing DSP binding to SERPINB5-GFP expressed in HaCaT sgNT1 cells. Expression of GFP served as negative control. **e-f)** Dispase-based dissociation assays with sgDPM1 HaCaT keratinocytes overexpressing SERPINB3, SERPINB4 and SERPINB5 (N=3). Each dot represents one biological replicate. One-way ANOVA, Dunnett’s multiple comparision test.

### SERPINB5 overexpression rescues desmoplakin localization and cytoskeletal distribution

Overexpression of SERPINB3, SERPINB4 and SERPINB5 in HaCaT keratinocytes showed no major biochemical changes in desmosomal protein expression (**Fig S3a, b**). However, SERPINB5 overexpression in DPM1 KO background led to rescue of DSP localization at cell junctions, indicating enhanced number of desmosomes under SERPINB5 rescue. SERPINB4, which showed no protective effect with regard to intercellular adhesion, did not ameliorate DSP membrane localization (**Fig 4a, b**). Similarly, SERPINB5 overexpression in primary human keratinocytes under DPM1 KO condition rescued DSP localization (**Fig 4c**). Consistent with these effects, SERPINB5 overexpression in DPM1 KO cells led to keratin organization similar to controls (**Fig 4d-f**), and a more regular distribution of the cortical actin belt underneath the plasma membrane (**Fig 4g-i**). These changes upon SERPINB5 rescue were also reflected in the mechanical properties of the cells, where SERPINB5 overexpression in HaCaT keratinocytes resulted in an increase in cell elasticity (**Fig 4j**). To address if SERPINB5 modulates DSP without the background of DPM1 loss, we targeted SERPINB5 by CRISPR/Cas9 (sgSERPINB5) in HaCaT keratinocytes (**Fig S3c**). Adhesion assays revealed significant loss of cell-cell adhesion in sgSERPINB5 cells (**Fig 4k, l**). In line with these observations, loss of SERPINB5 led to significantly reduced DSP localization at the cell membrane, indicating reduced desmosome numbers (**Fig 4m, n**). Thus, SERPINB5 modulates DSP membrane localization, even though the total protein levels of DSP remained unaltered (**Fig S3a, b**).

**Figure 4:**
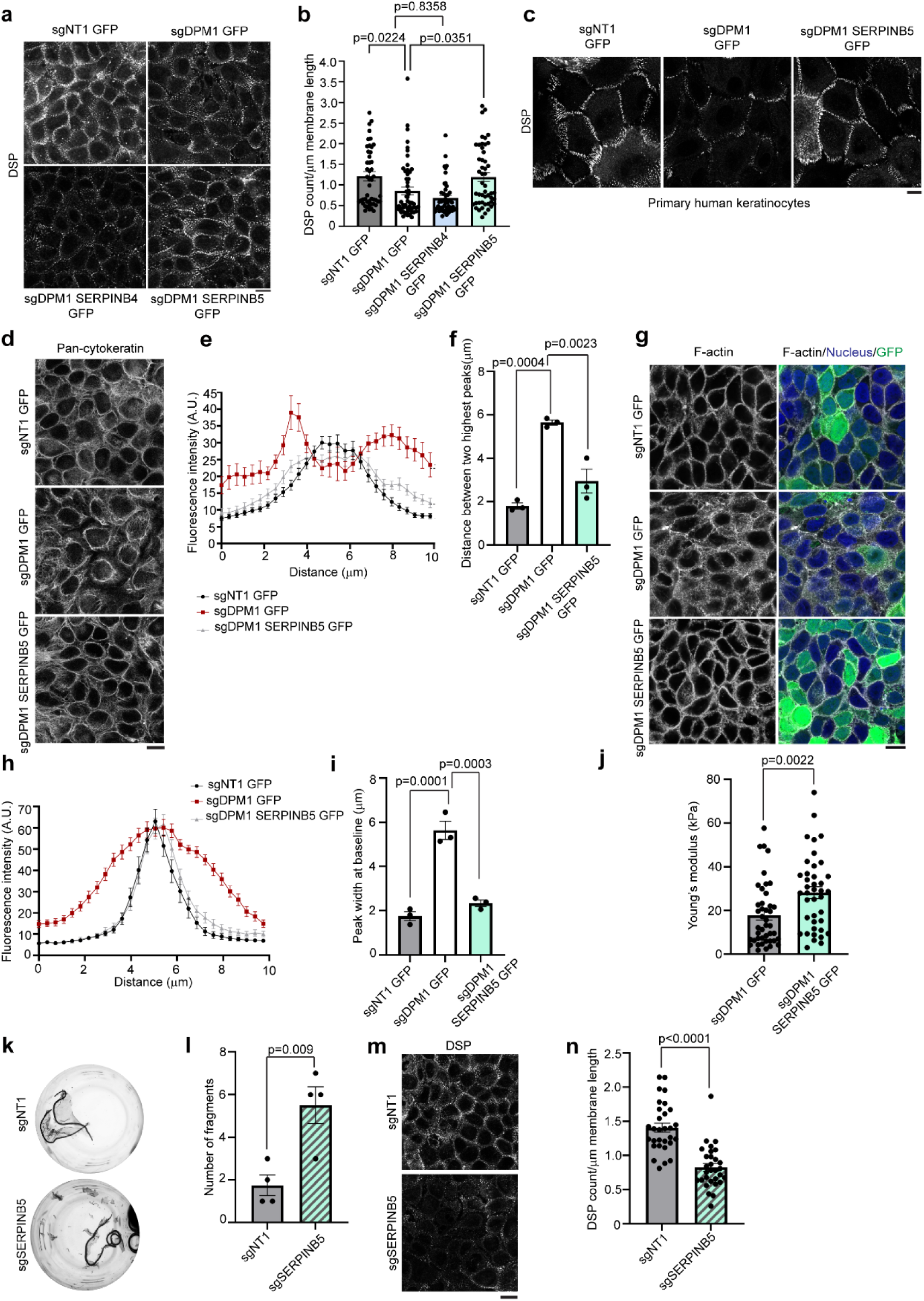
**a)** Images of DSP immunostainings in sgDPM1 and SERPIN overexpressing HaCaT keratinocytes. Panel shows representative of 3 biological replicates. Scale bar indicates 10µm distance. **b)** Quantification of the number of DSP puncta on the cell membrane normalised to the length of plasma membrane (µm). Each dot represents individual cells analysed from 3 independent biological replicates. One-way ANOVA, Dunnett’s multiple comparision test. **c)** Immunostaining showing DSP localization at the cell membrane upon SERPINB5 overexpression in DPM1 KO primary human keratinocytes. Panel shows representative of 3 biological replicates. Scale bar: 10µm distance. **d-f)** Keratin staining depicted by pan-cytokeratin in sgNT1, DPM1 KO and DPM1 KO overexpressing SERPINB5-GFP in HaCaT keratinocytes. Analysis done by measuring the keratin intensity across cell junctions (A.U.) over a distance of 10µm. Peak width calculated between the two highest points from the distribution plot profile graph. Scale bar: 10 µm distance. 30 individual cells were quantified from 3 independent biological replicates. One-way ANOVA, Dunnett’s multiple comparison test. **g-i)** F-actin staining by phalloidin in sgNT1, DPM1 KO and DPM1 KO overexpressing SERPINB5-GFP in HaCaT keratinocytes. Scale bar: 10 µm distance. Analysis done by measuring the phalloidin intensity across cell junctions (A.U.) over a distance of 10µm. The width of the peaks was calculated at baseline levels from distribution plot profile graphs. 30 individual cells were quantified from 3 independent biological replicates. One-way ANOVA, Dunnett’s multiple comparision test. **j)** Graph shows cellular elasticity, indicated by Young’s modulus (kPa). Each dot represents single cells from 3 biological replicates. Unpaired Student’s t-test. **k-l)** Dispase-based intercellular adhesion assays showing reduced cell-cell adhesion upon knockdown of SERPINB5 in HaCaT keratinocytes (N=4). Unpaired Student’s t-test. **m-n)** Images and analysis of DSP at the cell membrane upon loss of SERPINB5 (N=3). Each dot represents single cells from 3 biological replicates. Scale bar: 10 µm distance. Unpaired Student’s t-test.

### SERPINB5 prevents DPM1-induced differentiation defects in 3D-RHEs

Given the role of SERPINB5 in positively regulating cell-cell adhesion and DSP membrane localization in HaCaT and primary human keratinocytes, we asked if this would be reflected by changes in epidermal differentiation. Primary human keratinocytes transduced with sgDPM1 and either GFP or SERPINB5-GFP (**Fig 5a**) were allowed to differentiate for 12 days. Similar to **Fig 1f**, DPM1 KO epidermis expressing GFP showed a thickened cornified layer with intercellular gaps, indicating impairment of cell-cell adhesion. Upon SERPINB5 overexpression, the cornified layer was significantly thinner and comparable to the control conditions (**Fig 5b, c**). However, the thickness of the viable layers remained unchanged, resulting in a reduced total thickness of the epidermis (**Fig 5d, e**). In line with increased intercellular adhesion in response to SERPINB5 expression, the occurrence of intercellular gaps was also significantly diminished in SERPINB5-GFP expressing 3D-RHE (**Fig 5f**), while membrane localization of DSP on the cell surface was increased (**Fig 5g, h**). Interestingly, knockdown of DSP alone in 3D-RHEs, similar to DPM1 silencing, led to reduced thickness of non-corneal layers (**Fig 5k-m**), massive split formation (**Fig 5n**), and a disrupted staining pattern of the differentiation markers CK10 and filaggrin (**Fig 5j**). These data suggest that DPM1 together with SERPINB5 and at least in part by modulating DSP localization is required for epidermal differentiation.

**Figure 5:**
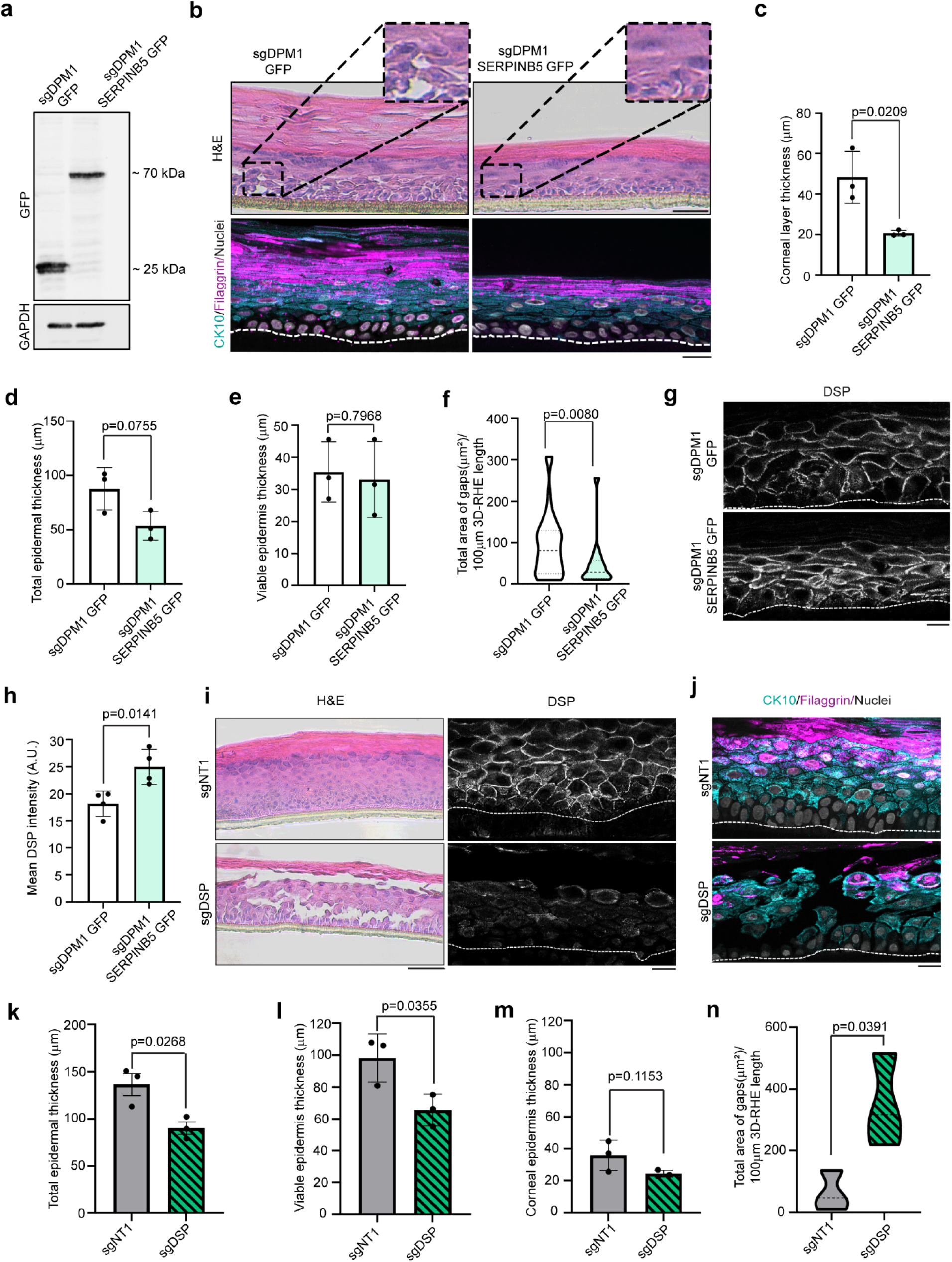
**a)** Western blot showing SERPINB5-GFP expression in primary human keratinocytes that lack DPM1. GFP served as control. GAPDH used as internal loading control. **b)** H&E staining of 3D-RHE from sgDPM1 expressing GFP control or SERPINB5-GFP 12 days post airlift. Immunostainings of CK10 and filaggrin were used as differentiation markers. Scale bar: 50µm distance on H&E images and 10µm on immunostaining images. White dashed line indicates insert membrane. Panel shows representatives from 3 biological replicates. **c-e)** Quantification of corneal layer thickness, total epidermal thickness and viable (non-corneal) epidermal layer thickness from 3 independent biological replicates. Student’s t-test, unpaired. **f)** Violin plot showing quantification of intercellular spaces within the epidermis in sgDPM1 GFP and sgDPM1 SERPINB5 GFP 3D-RHE. The area for defining intercellular spaces was set to be greater than 50µm^2^ to exclude shrinking artefacts. A minimum of 15 individual fields of view were used for analysis from 3 independent biological replicates. Unpaired Student’s t-test. **g-h)** Immunostaining and analysis of DSP intensity of sgDPM1 GFP and sgDPM1 SERPINB5-GFP 3D-RHEs. Panel shows representative of 4 biological replicates. Scale bar: 10µm distance. Unpaired Student’s t-test used to determine statistical significance (N=4). **i-j)** H&E staining of sgDSP 3D-RHEs 12 days post airlift. DSP staining shows depletion of DSP in sgDSP 3D-RHE. Immunostainings for CK10 and filaggrin used as differentiation markers. Scale bar: 50µm distance on H&E images and 10µm on immunostaining images. White dashed line indicates insert membrane. Panel shows representatives from 3 biological replicates. **k-m**) Quantification of total epidermal thickness, viable (non-corneal) epidermal layer thickness and corneal layer thickness from 3 independent biological replicates. Student’s t-test, unpaired. **n)** Violin plot showing quantification of intercellular spaces within the epidermis in sgNT1 and sgDSP 3D-RHE. A minimum of 15 individual fields of view were used for analysis from 3 independent biological replicates. Unpaired Student’s t-test.

### SERPINB5 and DPM1 modulate desmoplakin localization and cell-cell adhesion through S176 phosphorylation of DSP

It is known that the localization and cytoskeletal anchorage of DSP is regulated by phosphorylation ^7,28,29^. Given the role of DPM1 for DSP localization and keratin organization, we asked whether these changes are regulated by differential phosphorylation of DSP. Phospho-proteomics-based analysis of DSP revealed in total 18 sites spanning all domains of DSP to be phosphorylated in sgNT1 conditions (**Fig 6a**). Of these sites, only S176 and S2024 showed a significant and consistent change of phosphorylation in sgDPM1 cells with comparable total DSP levels (**Fig. 6a, Fig. S4a**). Interestingly, only the increase in S176 phosphorylation in sgDPM1 cells was prevented by SERPINB5 overexpression (**Fig 6b**), indicating SERPINB5-mediated modulation of this site. To elucidate the relevance of DSP phosphorylation at S176, we generated a phosphodeficient mutant (DSP-S176A) by exchange of serine to alanine (**Fig 6c, Fig S4b**). Interestingly, introduction of DSP-S176A in DPM1 KO cells led to significantly enhanced localization of DSP at the cell surface (**Fig 6c, d**) and the overall size of individual DSP complexes was enlarged in DSP-S176A cells. Further, while reconstitution of DSP-GFP in sgDPM1 cells was sufficient to improve cell-cell adhesion, this effect was stronger under expression of DSP-S176A (**Fig 6e, f**). These results suggest that SERPINB5 suppresses DSP phosphorylation at S176, which promotes DSP localization and intercellular adhesion at the cell membrane.

**Figure 6:**
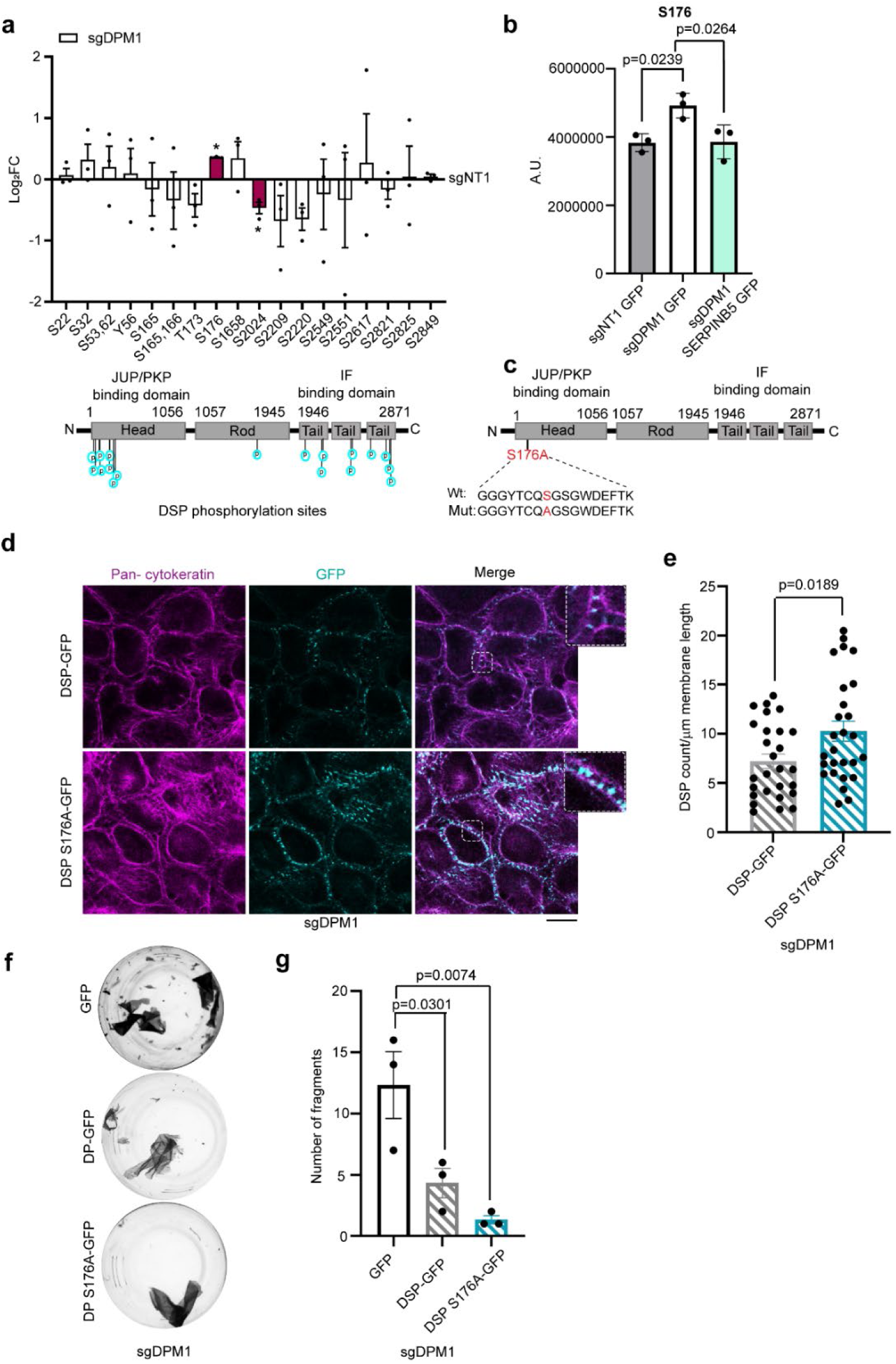
**a)** Graph showing differential phosphorylation sites of DSP in sgDPM1 HaCaT keratinocytes, presented as log_2_ fold change (FC) of respective values from sgNT1 control cells. Each dot represents one biological replicate. * indicates p<0.05, Student’s paired t-test. Red bars indicate significantly altered sites. Schematic of the representation of phospho-sites in DSP. **b)** Graph showing S176 phosphorylation intensity in sgNT1, sgDPM1 and sgDPM1 SERPINB5-GFP cells. Each dot represents one biological replicate. One-way ANOVA, Dunnett’s multiple comparision test (N=3). **c)** Schematic of the point mutation of S176 to A176 in the head domain of DSP. **d-e)** Images and analysis of DSP-GFP and DSP S176A-GFP expression at the cell membrane, in sgDPM1 HaCaT keratinocytes. Each dot represents individual cells from 3 biological replicates. Scale bar indicates 10 µm distance. Unpaired Student’s t-test (N=3). **f-g)** Dispase-based dissociation assays in sgDPM1 HaCaT keratinocytes upon reconstitution of DSP-GFP and DSP S176A GFP. Unpaired Student’s t-test (N=3). Each dot represents one biological replicate.

## DISCUSSION

### DPM1 and SERPINB5 may serve as link between differentiation and adhesion

The importance of O-linked and N-linked glycosylation in modulating cell-cell adhesion, desmosomes, epidermal differentiation and wound healing has been established in the past^14,15^. Since all glycosylation pathways are dependent on a basic mannose unit, we here used CRISPR/Cas9-based gene editing system to target DPM1, which is responsible for donating mannose to all important glycosylation steps. CRISPR/Cas9-based gene editing of DPM1 in a 3D organotypic model of human epidermis showed impaired stratification and differentiation, with reduced thickness of the viable epidermal layers and abnormally increased corneal layer thickness. Defects in differentiation were accompanied by intercellular gap formation in the epidermis, indicating loss of cell-cell adhesion. These results of DPM1 loss were prevented by overexpression of SERPINB5, suggesting that restoration of adhesion contributed to the rescue of differentiation defects. Indeed, despite the primary function in providing intercellular adhesion, desmosomal proteins are also known to modulate epidermal differentiation and signalling. As examples, loss of DSC1 and DSG4 led to differentiation defects in interfollicular epidermis^10,30^. It was also shown that DSG1 regulates epidermal differentiation, which is mechanistically dependent on EGFR and ERK signalling^31^. Ectopic expression of DSG3 in murine epidermis induces cornification defects and trans-epidermal water loss^32^ and DSG3 expression in suprabasal layers results in abnormal differentiation and hyperproliferation^33^. Further, the use of ectodomain-deleted DSG3 and inhibiting peptides led to impaired epidermal differentiation and epithelial morphogenesis^34,35^. These data all support a role of desmosomal adhesion in contributing to correct differentiation of the epidermis. Thus, the data are consistent with a role of DPM1, via SERPINB5, for epidermal homeostasis on the level of regulating intercellular adhesion. Although DPM1 is a crucial constituent of glycosylation pathways, it is so far unclear whether the changes we observed are mediated by altered glycosylation. Our data does not support a direct role of impaired glycosylation for membrane trafficking of adhesion molecules. However, indirect contributions, e.g., by altered glycosylation of other molecules which may be required for the stability and localization of desmosomal molecules, are of course possible. Further analyses will have to clarify the contribution of altered glycosylation in more detail.

### Mechanisms of DPM1-mediated modulation of cell-cell adhesion and differentiation

We found profoundly perturbed keratin and cortical actin organizations in DPM1 KO cells indicating that DPM1 modulates the complex interplay between the different components of the cytoskeleton and adhesion molecules. It is unclear in this context, whether reduced DSP at the cell surface leads to cytoskeletal remodelling or vice versa unstable DSP at cell junctions is a consequence of cytoskeletal impairments. A DSP mutant with higher binding affinity to keratin filaments strengthens intercellular adhesion^36^ and targeting DSG3 by autoantibodies from pemphigus patients induces uncoupling of keratins from the desmosome^37^. Vice versa, it has been demonstrated that loss of keratins from keratinocytes confer reduced binding forces and cell-cell junction mobility of DSG3^38^. Keratinocytes deficient for all keratins showed increased PKC-mediated phosphorylation of DSP, leading to destabilization of desmosomes^39^.The actin cytoskeleton, in contrast, appears to exert more indirect effects on desmosomal adhesion. It has been demonstrated that DSP assembly into desmosomes occurs in several stages with the last being actin-dependent^7^. Further, the actin-binding protein adducin and the cortical actin network are required for DSG3 membrane incorporation and cell-cell adhesion^40,41^. However, it has also been shown that desmosomal proteins in turn regulate actin organization in the cells^42,43^. Together, this profound interdependency explains the affection of both adhesion and cytoskeletal organization by DPM1 loss. Proteomic analysis of DPM1 KO keratinocytes revealed that cornification and cytoskeleton reorganization pathways were significantly upregulated, which is in line with the observed results discussed above.

However, DPM1 may regulate adhesion and differentiation also by mechanisms independent of the cytoskeleton and/or adhesion molecules. Metabolic pathways including carbohydrate metabolism and purine metabolism pathways were significantly downregulated under the loss of DPM1. Indeed, altered metabolic pathways have been directly associated with impaired epidermal differentiation. For example, altered lipid contents in the cornified layer is associated with skin conditions such as ichthyosis^44,45^. Detailed lipidomic analysis revealed specific lipid subsets which regulated epidermal differentiation and helped keratinocytes exit from the stem cell compartment^46^. Glucose metabolism is also important in maintaining skin homeostasis. mTOR pathway regulates glycolysis and mTOR effectors Rheb and Raptor KO in mice leads to differentiation defects in epidermis and reduced desmosomes^47^. In light of these data, it is likely that altered metabolic pathways contribute to the defects in epidermal differentiation in response to DPM1 loss.

### SERPINB5 mediates desmoplakin localization and intercellular adhesion

In an attempt to dissect the mechanisms underlying altered cell-cell adhesion in keratinocytes, we identified SERPINB5 as an interaction partner of interact DSP. BioID screening of the DSP interactome has shown the existence of SERPINB5 within the desmosomal plaque, supporting our findings^48^. The SERPIN class of proteins are classical serine peptidase inhibitors, however, SERPINB5 (MASPIN) is an exception. SERPINB5 does not have peptidase inhibitor activity^49^ and it’s detailed functions are largely unclear. It was studied majorly in the context of breast cancer where it modulates cell adhesion, migration and apoptosis of tumour cells^50–53^. SERPINB5 localization was shown to be regulated by EGFR and through cell density^54^, however how SERPINB5 modulates cell-cell adhesion and epidermal differentiation was not clear. We here show that deletion of SERPINB5 from HaCaT WT keratinocytes resulted in impaired DSP localization at the cell surface and loss of cell-cell adhesion, confirming that SERPINB5 is a regulator of DSP in cultured keratinocytes. Moreover, overexpressing SERPINB5 in DPM1 KO cells rescued DSP membrane localization and cytoskeletal alterations and reduced intercellular gap formation in 3D-RHEs. Corneal layer thickening in response to DPM1 loss was also ameliorated by ectopic expression of SERPINB5, although the thickness of the non-corneal epidermis was not affected. Nevertheless, this demonstrates an important contribution of SERPINB5 to the DPM1-dependent modulation of cornification. Along similar lines, SERPINB7 was recently shown to modulate epidermal differentiation and its loss led to psoriasiform lesions in mice^55^. These roles of the SERPINB family members in epidermal differentiation may warrant a more detailed analysis also with regard to conditions such as wound healing.

### DPM1 and SERPINB5 mediated alterations in DSP localization are partly modulated by phosphorylation of DSP

DSP localization at the cell membrane and association with keratin IF is known to be regulated by DSP phosphorylation. As example, dephosphorylation of S2849 in the tail domain of DSP leads to enhanced DSP-IF associations and strengthened cell-cell adhesion^29^. DSP phosphorylation at S2849 is regulated by a balance between GSK3 kinase and PP2A-B55a phosphatase, which regulates DSP localization and desmosome assembly in keratinocytes^29,56^. Our phospho-proteomics screen confirmed several known^28^ and identified some new phosphorylation sites in DSP, of which the phosphorylation of S176 was significantly increased in DPM1 KO cells and rescued upon SERPINB5 overexpression. Interestingly, dephosphorylation of an adjacent serine phosphorylation site, S165/S166, was recently shown to increase intercellular adhesion and keratin network integrity along with reduced phosphorylation of S2849^28^. As these sites were not significantly altered in DPM1 knockout cells, it appears that specifically S176 modulation is dependent on DPM1. As our data demonstrate that SERPINB5 directly interacts with DSP, it is possible that this interaction protects phosphorylation at S176. Understanding the kinase responsible for mediating this phosphorylation may add valuable insights into the understanding of the role of DSP phosphorylation for cell-cell adhesion and epidermal differentiation.

Taken together, we here showed that DPM1 by modulating SERPINB5 contributes to intercellular adhesion and epidermal differentiation. Specifically, DPM1 and SERPINB5 influence DSP membrane localization and the organization of both the keratin and cortical actin network. Finally, in this context, we identified a novel phosphorylation site of DSP (S176), which is modulated by SERPINB5 and suppressed under conditions of strong intercellular adhesion and correct DSP membrane localization (**Figure 7**).

**Figure 7:**
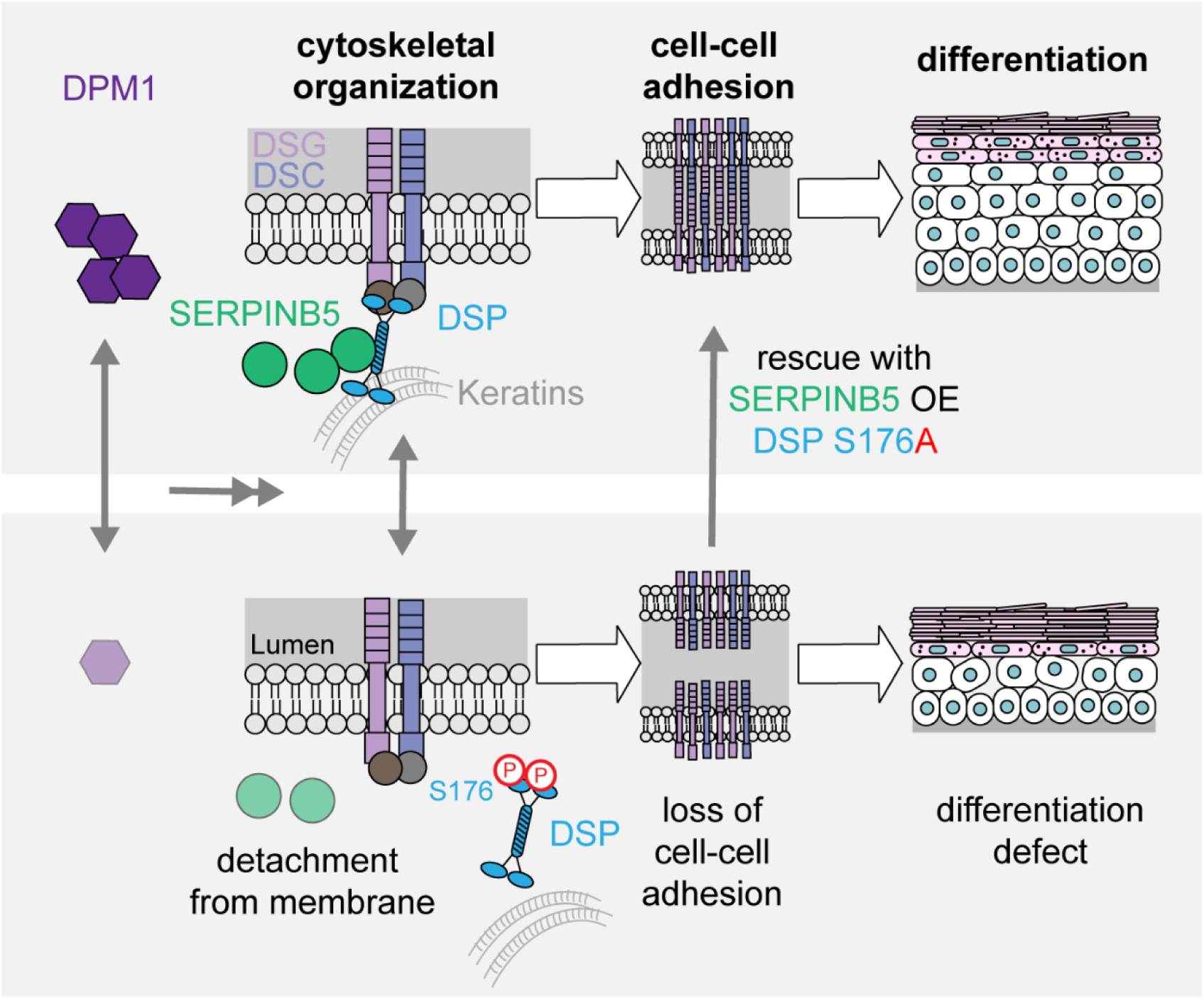
Schematic showing DPM1 regulates DSP expression on the cell membrane through SERPINB5, which can directly interact with DSP in keratinocytes. DPM1 loss has functional consequences on cell-cell adhesion, cytoskeletal organization and epidermal differentiation, which can be partly restored by rescue of SERPINB5 expression. Further, DSP localization on the surface is negatively regulated by phosphorylation of DSP on S176 position, which is significantly enhanced in DPM1 KO cells and interestingly SERPINB5 rescues the phosphorylation of DSP at S176 in DPM1 KO background.

## MATERIALS AND METHODS

### Cell culture and cloning

HaCaT keratinocytes^57^ were cultivated at 5% CO_2_ and 37°C in Dulbecco’s Modified Eagle Medium (DMEM) (Sigma-Aldrich, St. Louis, MO, USA) containing 1.8 mM calcium and complemented with 10% fetal bovine serum (Merck, Darmstadt, Germany), 50 U/ml penicillin, 50 µg/ml streptomycin (both AppliChem, Darmstadt, Germany) and 4 mM L-glutamine (Sigma-Aldrich). Primary human keratinocytes were cultured in EpiLife medium (#11684842 Fisher Scientific) containing 0.06 mM CaCl_2_ (Gibco, Carlsbad, California), supplemented with 1% Human Keratinocyte Growth Supplement (HKGS, #10761364 Fisher Scientific) and 1% Pen/Strep.

For cloning, oligos of sgDPM1, sgDPM2, sgDPM3, sgSERPINB5 and sgDSP were synthesized by microsynth (sequences mentioned in supplementary table 1), phosphorylated using T4 PNK enzyme (#0201S NEB, Ipswich, MA USA), annealed and ligated into Esp3I-digested (#R0734L NEB) lentiCRISPR_v2 (#52961 Addgene Watertown, USA) vector, using T4 ligase (#M0202S NEB). Ligated product was transformed in competent DH5α E. coli strain and plated on ampicillin-containing (100 ug/ml) agar plates. Colonies were amplified in 3 ml medium containing ampicillin (100 ug/ml). Plasmids were isolated via miniprep (#300287 Machery Nagel, Germany) and sequenced with U6_forward primer (GAGGGCCTATTTCCCATGATT).

For cloning SERPINB3, SERPINB4 and SERPINB5 overexpression constructs, PCR of HaCaT cDNA was performed using Platinum SuperFi 2 DNA Polymerase (#16410771 Fisher Scientific) according to manufacturer’s protocol and primers listed in table 1. The annealing temperature for the primers was set to 56°C. The PCR product was purified with PCR and gel purification kit (# 28506 Qiagen, Maryland, USA). The PCR product and the plasmid pLenti-C-mGFP (# PS100071 OriGene Technologies, Rockville, MD, USA) were digested with Asc1 (#R0558L Bioconcept, Allschwil, Switzerland) and Xho1 (#300366 NEB) for 3h at 37°C. The digested product was loaded on a 1% agarose gel and purified from the gel with PCR and gel purification kit (# 28506 Qiagen). The insert was ligated into the vector with a ratio of 3:1 overnight at 16°C using T4 Ligase (# 300361 Bioconcept). Positive clones were sequenced with forward and reverse primers (for TAATACGACTCACTATAGGG; rev CTTGATCTCCAGCTTGCCGT for SERPINB4; TCAGTGAAGCCAACACCAAGT for SERPINB3 and GAAAAGGAGCCACTGGGCAA for SERPINB5.

For generating DSP S176A mutant, Q5 Site-Directed Mutagenesis kit (#E0554S Bioconcept) was used and mutagenesis was performed as per standard kit instructions by the manufacturer. DSP-GFP^26^ plasmid was used as template. Annealing temperature was 63°C and extension time used was 7min (primer sequence in table 1). T7 forward primer (TAATACGACTCACTATAGGG) was used for sequencing.

### Generation of lentiviral constructs and stable cell lines

Lentiviral particles were generated according to standard procedures. HEK293T cells (between passage 9-11) were transfected with lentiviral packaging vector psPAX2 (#12259, Addgene, Watertown, MA, USA), the envelope vector pMD2.G (#12260, Addgene) and the respective construct plasmid using TurboFect (Thermo Fisher Scientific, Waltham, MA, USA). 48h post transfection, virus-containing supernatant was collected and concentrated using Lenti-Concentrator (OriGene), for minimum 2h at 4°C. Cells were transduced with the respective virus particles in an equal ratio using 5 µg/mL polybrene (Sigma-Aldrich) according to the manufacturer’s instructions. 24h post transduction for HaCaT keratinocytes and 8h later for primary human keratinocytes medium was exchanged and puromycin selection was given. Cells were cultivated for at least one week under selection, before starting with the respective experiments. Expression of the respective construct was confirmed via western blot analysis.

### Isolation of primary human keratinocytes

Foreskin tissue was obtained during circumcision of patients after informed consent according to the local ethical committee (EKNZ; date of approval: 11.06.2018, Project-ID: 2018-00963). Skin samples were washed three times in PBS containing 300 U/mL penicillin (#A1837 AppliChem), 300 U/mL streptomycin sulphate (#A1852 AppliChem) and 7.5 µg/mL amphotericin B (#A2942 Sigma-Aldrich). Excess tissue, blood vessels and parts of the dermis were removed and skin was cut into pieces of 0.5 x 1 cm size. To separate dermis and epidermis, skin samples were digested at 4°C overnight in 5 mg/mL dispase II solution (#D4693 Sigma-Aldrich) in HBSS (#H8264 Sigma-Aldrich) containing 300 U/mL penicillin, 300 U/mL streptomycin sulphate and 2.5 µg/mL amphotericin B. Epidermis was peeled off and washed once in PBS and digested in 0.25% trypsin and 1 mmol/L EDTA containing 100 U/mL penicillin and 100 U/mL streptomycin sulphate at 37°C for 20min. Trypsin activity was stopped by diluting 1+1 with a 1 mg/mL solution of soy bean trypsin inhibitor (#10684033 Gibco) in PBS. Keratinocytes were isolated by scratching the epidermis fragments on the dish bottom and through a 70 µm cell strainer (#431751 Corning, Somerville, USA). The isolated normal human epidermal keratinocytes (NHEK) were then seeded at a density of ~8 × 10^4^ cells/cm^2^ in EpiLife medium containing 60 µmol/L CaCl_2_ (#MEPI500CA Gibco) and 1% Human Keratinocyte Growth Supplement (#S0015 Gibco), 1% Pen/Strep and 2.5 µg/mL amphotericin B. After 3 days, the medium was exchanged, and from there on amphotericin B was omitted. For experiments, 40,000 cells were seeded in a 24-well plate and grown until they reached confluency. Subsequently, differentiation was induced by adding 1.2 mmol/L CaCl_2_ for 24h.

### 3D Reconstructed Human Epidermis (3D-RHE)

3D-RHE generation was performed according to the provided protocol of CellnTec (CellnTec, Bern, Switzerland). In brief, NHEK cells were cultivated in CnT-PR medium (#CNT-PR CellnTec) to ~50-70% confluency and transduced with respective constructs, as described above. Cells were selected in puromycin (Fisher Scientific) (2µg/ml). The cells were then seeded (1.7 x 10^5^ cells per insert) on 0.4 µm polycarbonate 24-well inserts (#10387523 Thermo Scientific) which were coated with 15 µg/cm^2^ rat-tail collagen I (#50201-IBIDI, Gräfelfing, Germany) in 0.02 M acetic acid at 37°C for 1h. NHEKs were cultivated in CnT-PR medium. 24h after seeding, the medium inside the insert and 6-well was exchanged with CnT-PR-3D medium (#CnT-PR-3D CellnTec). After 16-8h, inserts were lifted to an air-liquid interface by aspirating the media inside the insert and adding CnT-PR-3D in the 6-well up to the level of the insert membrane (1.3 mL); the CnT-PR-3D medium was exchanged every other day. The 3D organotypic cultures were harvested and analyzed 12 days post-airlifting by fixing in 2% paraformaldehyde at 4°C for 4h.

### Western Blot

Confluent cell monolayers were lysed with SDS lysis buffer (25 mM HEPES, 2 mM EDTA, 25 mM NaF, 1% SDS, pH 7.6) supplemented with an equal volume of a protease inhibitor cocktail (cOmplete, Roche Diagnostics, Mannheim, Germany), by using a cell scraper. Lysates were sonicated and the total protein amount was determined with a BCA protein assay kit (Thermo Fisher Scientific) according to the manufacturer’s instructions. The proteins were denatured by heating in Laemmli buffer, for 10min at 95°C. Membranes were blocked in odyssey blocking buffer (Li-Cor, Lincoln, NE, USA) for 1h at room temperature. The following primary antibodies were diluted with odyssey blocking buffer in tris-buffered-saline containing 0.1% Tween20 (TBS-T) (Thermo Fisher Scientific) and incubated overnight at 4°C, with rotation: mouse Dsc2 mAb (#60239-1-Ig Proteintech, Rosemont, IL, USA), mouse Dsg2 mAb (clone 10G11, #BM5016 Acris, Herford, Germany), rabbit Dsg3 pAb (EAP3816 Elabscience, Biozol, Eching, Germany), mouse Pkp1 mAb (clone 10B2, #sc-33636 Santa Cruz, Dallas, TX, USA), mouse Pkp2 mAb (#651101 Progen, Heidelberg, Germany), mouse PG mAb (clone PG5.1, #61005 Progen), mouse DSP mAb (#sc-390975 Santa Cruz), mouse E-Cad mAb (clone 36, #610181 BD Biosciences, Franklin lakes, NJ, USA), rabbit DSC1 mAb (#ab150382 Abcam, Cambridge, UK), mouse DSG1 ((#651111 Progen), rabbit keratin 10 (#905403 Biolegend), mouse keratin 14 (ab7800 Abcam), rabbit DPM1 (#12403-2-AP, Proteintech), rabbit SERPINB5 (MASPIN #ab182785 Abcam), rabbit mGFP (#TA150122 Thermo Fisher Scientific), mouse GAPDH mAb (clone 0411, #sc-47724 Santa Cruz), mouse α-tubulin (clone 10D8, #627901 BioLegend). Goat anti-mouse 800CW and goat anti-rabbit 680RD (#925-32210 and #925-68071, both Li-Cor) were used as secondary antibodies, incubated for 1h at room temperature. Odyssey FC imaging system was used for imaging the blots and band density was quantified with Image Studio (both Li-Cor).

### Immunoprecipitation

Confluent cell monolayers were washed twice with ice-cold PBS and incubated with modified RIPA buffer plus protease inhibitors (10 mM Na_2_HPO_4_, 150 mM NaCl, 1%Triton X-100, 0.25% SDS, 1% sodium deoxycholate, pH 7.3), for 30min on ice. Cells were then scraped and homogenized on ice by passing 10 times through a 20G and 25G injection needle. Cell debris were removed by centrifugation at 7000g, 5min, 4°C. Protein concentration was determined by BCA (Thermo Fisher Scientific) and equal amounts of protein were diluted in 1ml of RIPA buffer. GFP trap magnetic agarose beads (gtma-20 ChromoTek, Martinsried, Germany) were washed 3 times with 1ml RIPA buffer and 25µl of beads were added to diluted lysates and incubated for 1.5h at 4°C. Beads were washed twelve times with wash buffers and afterwards mixed with Laemmli buffer and denaturized at 95°C for 10min. The following samples from I.P were subjected to western blotting as described above. Rabbit mGFP (#TA150122 Thermo Fisher Scientific) primary antibody was used to confirm immunoprecipitation and mouse DSP mAb (#sc-390975 Santa Cruz) antibody was used to detect DSP co-immunoprecipitation.

### Dispase - based dissociation assay

HaCaT keratinocytes expressing various sgRNA and overexpression constructs or primary human keratinocytes were seeded in equal numbers in 24-well plates. After reaching confluency, cells were washed with pre-warmed PBS and incubated with 250 µl dispase II (50mg/10ml in HBSS; Sigma-Aldrich D4693) for 20min (HaCaTs) and 45min for primary human keratinocytes at 37°C to detach the intact cell sheet from the well bottom. 150 ul HBSS was added to these wells and a constant mechanical shear stress was applied using an electrical pipette (Eppendorf, Hamburg, Germany), for 10 times each well. Wells were finally imaged with a binocular microscope (Olympus, Tokyo, Japan) and SLR camera (Canon, Tokyo, Japan). The number of fragments generated are a direct measure of cell cohesion, which is inversely proportional to adhesive strength.

### Immunostaining

Cells were grown on 13 mm glass coverslips and fixed with either 2% PFA (Thermo Fisher Scientific) in PBS at room temperature or ice-cold methanol (Merck Millipore) for 10min on ice. Cells were permeabilized with 0.1% Triton X-100 in PBS for 10min and blocked with 3% BSA and 1% normal goat serum in PBS for 1h, in a humidified chamber. The following primary antibodies were incubated overnight at 4°C: mouse Dsg2 mAb (clone 10G11, #BM5016 Acris), rabbit Dsg2 pAb (#610121 Progen), mouse Dsg3 mAb (clone 5G11, # 326300 Invitrogen, Carlsbad, CA, USA), rabbit DSP pAb (NW39), mouse DSP mAb (1G4) (both kind gifts from Kathleen Green, Northwestern University, Chicago, USA), Phalloidin CruzFluor 488 (#sc-363791 Santa Cruz), pan-cytokeratin (AE1/AE3) efluor 570 (#41-9003-80 eBioscience). Following primary antibody incubation, cells were washed 3 times with PBS and AlexaFluor (AF488, AF568) conjugated anti-rabbit or anti-mouse antibodies (#A-11008, A-11004 Fisher Scientific) were added and incubated for 1h at room temperature (RT). DAPI (Sigma-Aldrich) was added for 10min to counterstain nuclei. Cells were washed 3 times with PBS and mounted with ProLong Diamond Antifade (Thermo Fisher Scientific). Image acquisitions were done using Stellaris 8 Falcon confocal microscope (Leica, Wetzlar, Germany) with a HC PL APO CS2 63x/1.40 oil objective. Image analysis was done with ImageJ software for analyzing plot profiles and counting DSP puncta as described later in data analysis section.

For staining of paraffin sections, tissues embedded in paraffin were cut into 5 µm thick sections with an automated microtome (HM355S, Thermo Fisher Scientific). After deparaffinization, temperature-mediated antigen retrieval was performed in citrate buffer (10 mM citric acid monohydrate (VWR, 20276.235, Pennsylvania, USA), Ph 6, 0.1% triton X-100) for 20min at 95°C. Tissue was permeabilized in 0.1% triton X-100 in PBS for 5min and blocked with 3% bovine serum albumin/0.12% normal goat serum in PBS for 1h. The following primary antibodies were incubated in PBS at 4°C overnight: mouse DSP (#61003 Progen), rabbit DSG2 RB5 (# 610120 Progen Biotechnik), Filaggrin (#ab218395 Abcam), Keratin 10 (#905403 Biolegend), Ki67 B56 (#550609 BD Bioscience), rabbit SERPINB5 (MASPIN #ab182785 Abcam), rabbit DPM1 (#12403-2-AP, Proteintech). Respective secondary goat anti-rabbit or goat anti-mouse antibodies coupled to Alexa Fluor 488, Alexa Fluor 568 (both Thermo Fisher Scientific), or cy5 (Dianova, Hamburg, Germany) were incubated for 1h at RT and DAPI (Sigma-Aldrich) was added for 10min to counterstain nuclei. Finally, samples were mounted with Fluoromount Aqueous Mounting Medium (Sigma-Aldrich).

### Histology

Fixed 3D-RHEs were removed from the cell culture insert using a 8 mm biopsy punch (#600213 Stiefel), cut into half and embedded in histogel (#HG-4000-012 Epredia). After polymerization of the histogel, samples were paraffin embedded using the TPC 15 Tissue Processor (Medite Medizintechnik): 1) 70% ethanol, 37°C, 45min; 2) 80% ethanol, 37°C, 45min; 3) 96% ethanol, 37°C, 30min; 4) 96% ethanol, 37°C, 45min; 5) 100% ethanol, 37°C, 30min; 6) 100% ethanol, 37°C, 60min; 7) 100% ethanol, 37°C, 60min; 8) Xylene, 37°C, 30min; 9) Xylene, 37°C, 45min; 10) Xylene, 37°C, 60min; 11) Paraffin, 62°C, 45min; 12) Paraffin, 62°C, 60min; 13) Paraffin, 62°C, 60min. TES Valida embedding station (Medite Medizintechnik) was used to embed processed tissue into paraffin blocks and cut into 5 µm sections using an automated microtome (HM355S, Thermo Fisher Scientific). Haematoxylin and Eosin (H&E) staining was performed according to standard procedures. In brief, sections were stained with mayer’s haemalaun solution (#1.09249.1022 Sigma-Aldrich) for 5min, washed, dehydrated in an increasing ethanol series and stained with 0.5% (w/v) eosin solution for 5min. After washing steps in ethanol and methyl salicylate, sections were mounted with DPX mounting media (#06522 Sigma-Aldrich).

### Atomic Force Microscopy (AFM)

A Nanowizard IV atomic force microscope (AFM, JPK Instruments, Berlin, Germany) mounted on an inverted fluorescence microscope (IX83, Olympus) was used for cell stiffness measurements. Experiments were performed in culture medium at 37°C using Si_3_N_4_ AFM cantilevers (pyramid-shaped D-tip, MLCT cantilever, Bruker, 4 Billerica, MA, USA) with a spring constant of 0.03 N/m. Spring constant was calibrated for each cantilever at 37°C applying the thermal noise method. Force-displacement curves were obtained in force spectroscopy mode using the following settings: relative setpoint 0.4 nN, z-length 5 μm, extend delay 0s, pulling speed 2 μm/s, and recorded with the SPM control v.4 software 15 (JPK Instruments). Cells were seeded on glass coverslips. All measurements were performed on the cell center and conducted within 1h. Force distance curves were analyzed using JPKSPM Data Processing software (version 6, JPK Instruments) and then fitted with the Hertz model for young’s modulus calculation. The tip-half opening angle was 17.5 degrees, the Poisson’s ratio was set as 0.5 and 300 nm of indentation depth was used.

### Fluorescence Recovery After Photobleaching (FRAP)

For FRAP measurements, HaCaT cells (sgNT1) and sgDPM1 were transduced with DSP-GFP^26^ as described above and seeded in 8-well imaging chambers (IBIDI). After the formation of visible junctions, FRAP measurements were performed on a Stellaris 8 Falcon confocal microscope (Leica) with a HC PL APO CS2 63x/1.40 oil objective, at 37°C with 5% CO_2_ and constant humidity. The measurements were carried out and analyzed with the fluorescence recovery after photobleaching wizard software (Leica). Regions of interests were defined along cell-cell junctions containing a desmosome between two neighboring mGFP positive cells. After 5 frames of recording the prebleach intensity, mGFP signal was bleached shortly for 5 frames, using the 488 nm laser line at 80% transmission on FRAP booster mode and the fluorescence recovery was recorded over 239s with 100 frames for the initial 26s and 45 frames for the remaining time. The fraction of mobile molecules was determined by the formula: Mobile fraction = Ie – Io/ Ii – Io, where, Ie is the intensity reached after recovery time, Io is the minimal intensity that was achieved right after bleaching, and Ii is the average prebleach intensity value.

### Proteomics

#### Sample preparation for enrichment analysis

Samples were adjusted to the 5% of SDS, 100 mM TEAB and 10 mM TCEP and subsequently reduced for 10min at 95°C. Samples were then cooled down to RT and 0.5 µL of 1M iodoacetamide was added to the samples. Cysteine residues were alkylated for 30min at 25°C in the dark. Digestion and peptide purification were performed using S-trap technology (Protifi) according to the manufacturer’s instructions. In brief, samples were acidified by addition of 2.5 µL of 12% phosphoric acid (1:10) and then 165 µL of S-trap buffer (90% methanol, 100 mM TEAB pH 7.1) was added to the samples (6:1). Samples were briefly vortexed and loaded onto S-trap micro spin-columns (Protifi) and centrifuged for 1 min at 4000g. Flow-through was discarded and spin-columns were then washed 3 times with 150 µL of S-trap buffer (each time samples were centrifuged for 1 min at 4000 g and flow-through was removed). S-trap columns were then moved to the clean tubes and 20 µL of digestion buffer (50 mM TEAB pH 8.0) and trypsin (at 1:25 enzyme to protein ratio) were added to the samples. Digestion was allowed to proceed for 1h at 47°C. After, 40 µL of digestion buffer was added to the samples and the peptides were collected by centrifugation at 4000g for 1min. To increase the recovery, S-trap columns were washed with 40 µL of 0.2% formic acid in water (400g, 1min) and 35 µL of 0.2% formic acid in 50% acetonitrile. Eluted peptides were dried under vacuum and stored at −20°C until further analysis.

#### Data acquisition for enrichment analysis

Dried peptides were resuspended 0.1% FA and 0.2 ug of peptides were subjected to LC–MS/MS analysis using a Q Exactive Plus Mass Spectrometer fitted with an EASY-nLC 1000 (both Thermo Fisher Scientific) and a custom-made column heater set to 60°C. Peptides were resolved using a RP-HPLC column (75μm × 30cm) packed in-house with C18 resin (ReproSil-Pur C18–AQ, 1.9 μm resin; Dr. Maisch GmbH) at a flow rate of 0.2 ml/min-1. A linear gradient ranging from 5% buffer B to 45% buffer B over 60min was used for peptide separation. Buffer A was 0.1% formic acid in water and buffer B was 80% acetonitrile, 0.1% formic acid in water. The mass spectrometer was operated in DDA mode with a total cycle time of approximately 1s. Each MS1 scan was followed by high-collision-dissociation (HCD) of the 20 most abundant precursor ions with dynamic exclusion set to 5s. For MS1, 3e^6^ ions were accumulated in the Orbitrap over a maximum time of 25ms and scanned at a resolution of 70,000 FWHM (at 200 m/z). MS2 scans were acquired at a target setting of 1e^5^ ions, maximum accumulation time of 110ms and a resolution of 35,000 FWHM (at 200 m/z). Singly charged ions, ions with charge state ≥ 6 and ions with unassigned charge state were excluded from triggering MS2 events. The normalized collision energy was set to 27%, the mass isolation window was set to 1.4 m/z and one microscan was acquired for each spectrum.

#### Sample preparation for total proteomics

Cells were lysed in 50 µL of lysis buffer (1% Sodium deoxycholate (SDC), 10 mM TCEP, 100 mM Tris, pH=8.5) using twenty cycles of sonication (30s on, 30s off per cycle) on a Bioruptor (Dianode). Following sonication, proteins in the bacterial lysate were reduced by TCEP at 95°C for 10min. Proteins were alkylated using 15 mM chloroacetamide at 37° C for 30min and further digested using sequencing-grade modified trypsin (1/50 w/w, ratio trypsin/protein; Promega, USA) at 37°C for 12h. After digestion, the samples were acidified using TFA (final 1%). Peptide desalting was performed using iST cartridges (PreOmics, Germany) following the manufactures instructions. After drying the samples under vacuum, peptides were stored at −20°C and dissolved in 0.1% aqueous formic acid solution at a concentration of 0.5 mg/ml upon use.

#### Data acquisition for total proteomics

Dried peptides were resuspended 0.1% FA and 0.2 ug of peptides were subjected to LC–MS/MS analysis using a Orbitrap Fusion Lumos Mass Spectrometer fitted with an EASY-nLC 1200 (both Thermo Fisher Scientific) and a custom-made column heater set to 60°C. Peptides were resolved using a RP-HPLC column (75μm × 36cm) packed in-house with C18 resin (ReproSil-Pur C18–AQ, 1.9 μm resin; Dr. Maisch GmbH) at a flow rate of 0.2 μL/min-1. The following gradient was used for peptide separation: from 5% B to 12% B over 5min to 35% B over 65min to 50% B over 20min to 95% B over 2min followed by 18min at 95% B. Buffer A was 0.1% formic acid in water and buffer B was 80% acetonitrile, 0.1% formic acid in water. The mass spectrometer was operated in DDA mode with a cycle time of 3s between master scans. Each master scan was acquired in the Orbitrap at a resolution of 120,000 FWHM (at 200 m/z) and a scan range from 375 to 1500 m/z followed by MS2 scans of the most intense precursors in the linear ion trap at “Rapid” scan rate with isolation width of the quadrupole set to 1.4 m/z. Maximum ion injection time was set to 50ms (MS1) and 35ms (MS2) with an AGC target set to 1e^6^ and 1e^4^, respectively. Only peptides with charge state 2 – 5 were included in the analysis. Monoisotopic precursor selection (MIPS) was set to Peptide, and the Intensity Threshold was set to 5e^3^. Peptides were fragmented by HCD (Higher-energy collisional dissociation) with collision energy set to 35%, and one microscan was acquired for each spectrum. The dynamic exclusion duration was set to 30s.

#### Phospho proteomics

Cells were lysed in 2M guanidium hydrochloride, 100 mM ammonium bicarbonate, 5mM TCEP, phosphatase inhibitors (Sigma P5726&P0044) by sonication (Bioruptor, 10 cycles, 30s on/off, Diagenode, Belgium). Proteins were subsequently reduced by 10min incubation at 95°C and alkylated with 10 mM chloroacetamide for 30min at 37 °C. Samples were then diluted with 100 mM ammonium bicarbonate to a final guanidium hydrochloride concentration of 0.5 M. Proteins were digested by incubation with sequencing-grade modified trypsin (1/50, w/w; Promega, Madison, Wisconsin) for 12h at 37°C. After digestion, samples were acidified with 5% TFA and peptides were purified using C18 reverse-phase spin columns (Macrospin, Harvard Apparatus) according to the manufacturer’s instructions, dried under vacuum and stored at −20°C until further use. Peptide samples were enriched for phosphorylated peptides using Fe(III)-IMAC cartridges on an AssayMAP Bravo platform. Phospho-enriched peptides were resuspended in 0.1% aqueous formic acid and subjected to LC–MS/MS analysis using a Orbitrap Fusion Lumos Mass Spectrometer fitted with an EASY-nLC 1200 (both Thermo Fisher Scientific) and a custom-made column heater set to 60°C. Peptides were resolved using a RP-HPLC column (75μm × 36cm) packed in-house with C18 resin (ReproSil-Pur C18–AQ, 1.9 μm resin; Dr. Maisch GmbH) at a flow rate of 0.2 μLmin-1. The following gradient was used for peptide separation: from 5% B to 8% B over 5min to 20% B over 45min to 25% B over 15min to 30% B over 10min to 35% B over 7min to 42% B over 5min to 50% B over 3min to 95% B over 2min followed by 18min at 95% B. Buffer A was 0.1% formic acid in water and buffer B was 80% acetonitrile, 0.1% formic acid in water. The mass spectrometer was operated in DDA mode with a cycle time of 3s between master scans. Each master scan was acquired in the Orbitrap at a resolution of 120,000 FWHM (at 200 m/z) and a scan range from 375 to 1600 m/z followed by MS2 scans of the most intense precursors in the Orbitrap at a resolution of 30,000 FWHM (at 200 m/z) with isolation width of the quadrupole set to 1.4 m/z. Maximum ion injection time was set to 50ms (MS1) and 54 ms (MS2) with an AGC target set to 250% and “Standard”, respectively. Only peptides with charge state 2 – 5 were included in the analysis. Monoisotopic precursor selection (MIPS) was set to Peptide, and the Intensity Threshold was set to 2.5e^4^. Peptides were fragmented by HCD (Higher-energy collisional dissociation) with collision energy set to 30%, and one microscan was acquired for each spectrum. The dynamic exclusion duration was 30s.

### Image Processing and Statistics

Figures were compiled with Photoshop CC and Illustrator CC (Adobe, San José, CA, USA). Statistical analysis was performed using GraphPad Prism 8 with a two-tailed student’s t-test for the comparison of two data sets and one-way or two-way ANOVA corrected by either Dunnett’s, Dunn’s or Tukey’s test for more than two data sets. Statistical significance was determined at p < 0.05. The data sets were first tested for normal distribution using Shapiro-Wilk normality test and respective statistics were applied based on the distribution. Welch correction for unequal variances was applied where applicable. The error bars shown in all graphs were depicted as ± SD. A minimum of 3 biological replicates was used in each experiment.

### Data Analysis

#### Epidermal thickness analysis

Epidermal thickness was analyzed on H&E-stained tissue sections by using the image analysis software QuPath^58^. In brief, 3-5 images per section were analyzed in QuPath by marking a rectangular area of the tissue section and then applying the “wand” tool to mark the different layers of the epidermis. The area of these were measured and divided by the total length of the epidermis. The following formula was applied to calculate the tissue thickness.

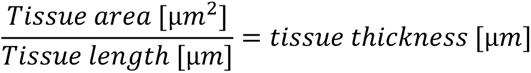

Corneal layer and non-corneal layer (viable) thickness were normalized to total epidermal thickness, to account for variances in overall thickness across biological replicates.

#### Ki67 analysis of 3D-RHE

Proliferation of 3D-RHE was analyzed on Ki67 staining from tissue sections, using QuPath software^58^. In brief, 3-5 images per sample were analyzed in QuPath and nuclei were masked by detecting DAPI signal. The total numbers of nuclei per field were quantified. From these masks generated, Ki67 positive nuclei were counted and the ratio of Ki67 positive nuclei to total nuclei was calculated to determine the percentage of Ki67 positive cells.

#### ImageJ Analysis

Image J was used to generate kymographs of FRAP recovery data for DSP GFP signal over dynamic time scales before and after bleaching.

Keratin distribution and cortical actin distribution were analyzed by drawing a ROI rectangle of fixed size at the cell-cell contact sites (10 micron) and measuring the mean plot profile intensities along this area for individual cells. GraphPad PRISM software was used to create graphs and statistical analysis.

DSP dots were quantified by applying a threshold to the DSP staining on images, such that all the DSP puncta were accounted for. The membrane length for individual cells was measured by drawing ROI around the cell membranes and the respective number of DSP puncta over this cell were counted using particle analysis in ImageJ.

#### Proteomic data analysis

The raw data were analyzed using MaxQuant (v1.6.17.0)^59^ with default setting. In brief, the spectra were searched against a human database (protein sequences downloaded from www.uniprot.org on 2020/04/17) and commonly observed contaminants by the Andromeda search engine ^60^, using the following search criteria: full tryptic specificity was required (cleavage after lysine or arginine residues, unless followed by proline); 3 missed cleavages were allowed; carbamidomethylation (C) was set as fixed modification; oxidation (M) and acetylation (Protein N-term) were applied as variable modifications; mass tolerance of 20 ppm (precursor) and 20 ppm/0.6 Da (fragments) for QE-plus/Lumos. Label-free and iBAQ quantification as well as much between runs were enabled. The database search results were filtered to a false discovery rate (FDR) to 1% on the peptide and protein level and specified a minimum length of 7 amino acids for peptides. Quantitative analysis results from label-free quantification were processed using the MSstats R package v4.0.1^61^.

Heat maps were generated by using ClustVis software freely available as an online tool. STRING database was used to identify molecular and biological pathways associated with the significantly modulated proteins across the samples.

## ACKNOWLEDGEMENTS

The authors would like to acknowledge Dr. Diego Calabrese (Histology Core Facility); and Drs. Michael Abanto and Pascal Lorentz (Microscopy Core Facility), University of Basel, Switzerland, Dr. Florian Geier, Bioinformatics Facility Basel (University of Basel), Prof. Kathleen Green, Northwestern University, USA) for providing desmoplakin antibodies and Addgene plasmid #32227, Department of Urology (University hospital, Basel) for providing human foreskin tissues, Swiss National Science Foundation-SNSF (#197764 to Prof. Dr. Spindler), Swiss Heart Foundation (FF21098 to CS and VS), the Olga Mayenfisch Stiftung and the Novartis Foundation for medical-biological research (#22B086 both to CS) for funding.

## AUTHOR CONTRIBUTIONS

Conceptualization: VS, MR; Data acquisition: MR, HF, VB, MTW, KLF, PH, CST, AZ, KB; Data Analysis: MR, VB, MTW, KLF, KB; Funding Acquisition: VS, CS; Project Administration: VS, MR; Resources: VS, CS; Supervision: VS; Writing Original Draft Preparation: MR, VS; Writing - Review and Editing: VS, MR, HF, VB, MTW, KLF, PH, CST, AZ, CS.

