## Supplementary for "DPM1 through SERPINB5 modulates desmosomal adhesion and epidermal differentiation"

**Supplementary Figures:**

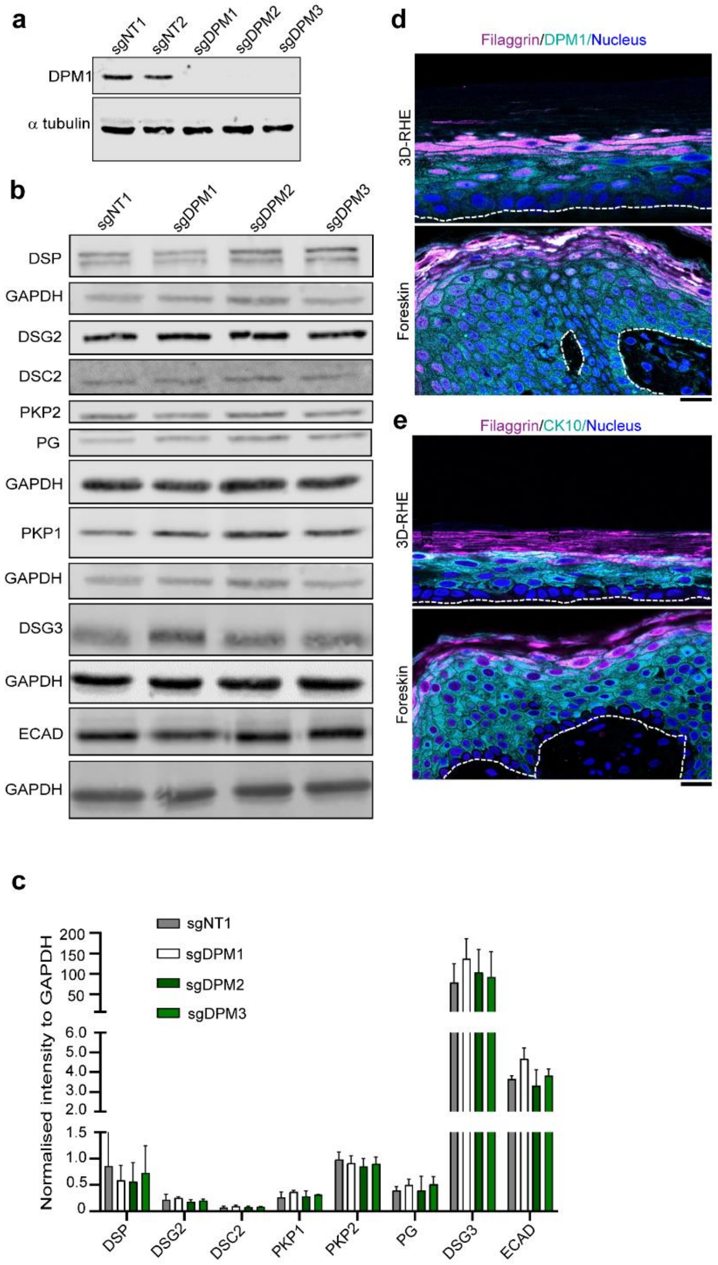


**Figure S1: a)** Western blot showing DPM1 in sgDPM1, sgDPM2 and sgDPM3 HaCaT keratinocytes. α tubulin was used as internal loading control. **b-c)** Western blot images and quantifications of desmosomal proteins from sgDPM1, sgDPM2 and sgDPM3 HaCaT keratinocytes. Representative images of 3 biological replicates are shown. GAPDH was used as internal loading control (N=3) **d)** Immunofluorescence staining of DPM1 and filaggrin in 3D-RHE and human foreskin tissue, as indicated. Filaggrin was used as a differentiation marker. Dashed line indicates insert membrane /basement membrane. Scale bar: 10µm. Panel shows representative of 3 biological replicates **e)** Immunostaining of CK10 and filaggrin in 3D-RHE and human foreskin tissue, as indicated. Dashed line indicates insert and basement membrane. Scale bar: 10µm distance. Panel shows representative of 3 biological replicates.

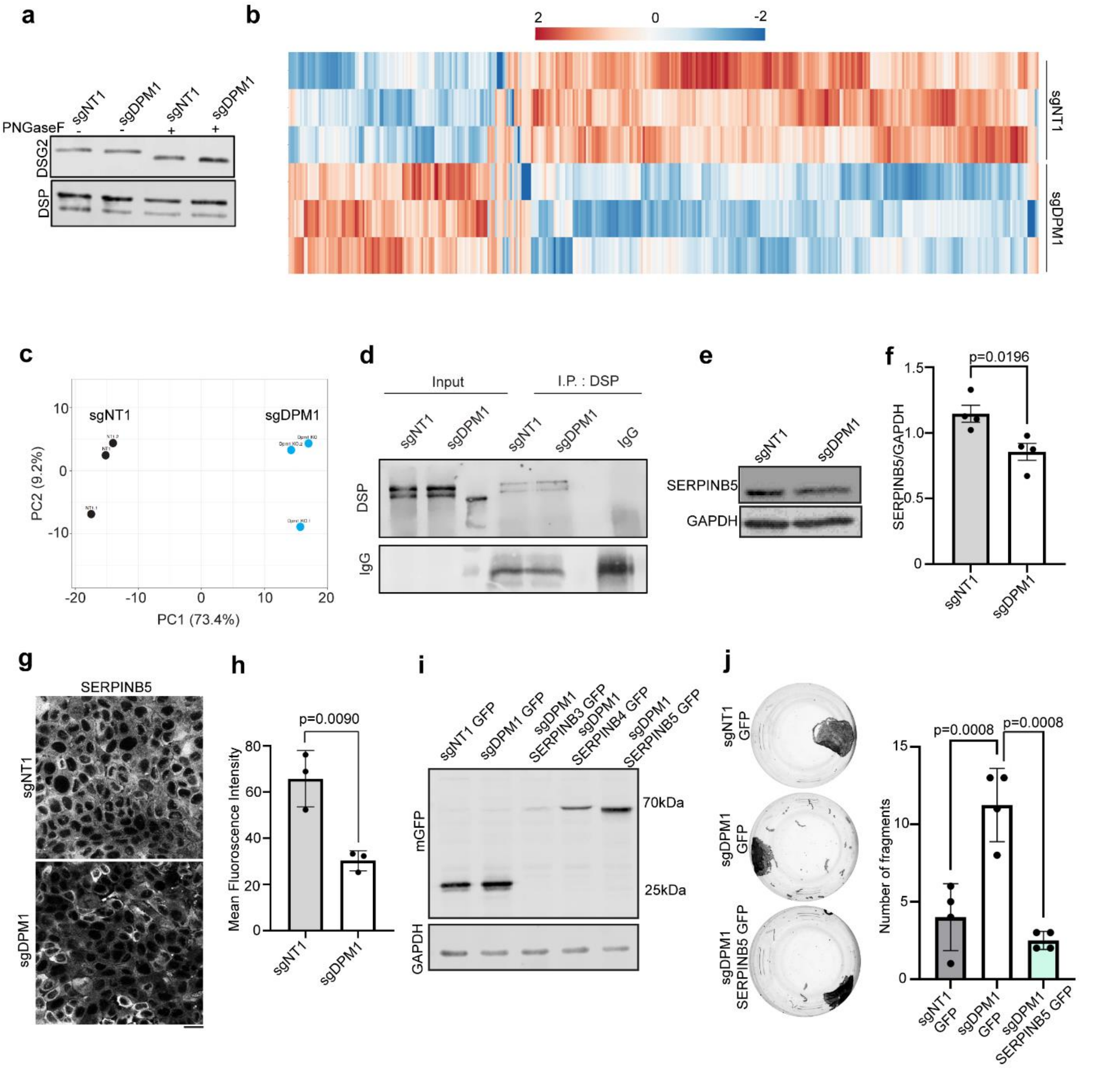


**Figure S2: a)** Western blot showing migration of DSG2 and DSP upon treatment with PNGasF in sgNT1 versus sgDPM1 HaCaT keratinocytes. **b)** Heat map showing differential protein expression profiles in sgNT1 and sgDPM1 HaCaT keratinocytes from 3 biological replicates. Color scale represents $\log_2$ fold change of intensity values **c)** Principle component (PC) analysis of the samples used in b) **d)** Immunoprecipitation assay showing DSP pulldown in sgNT1 and sgDPM1 cells. IgG used as negative control. Panel shows representative of 2 biological replicates. **e-f)** Western blot and corresponding quantification of SERPINB5 expression in sgDPM1 HaCaT keratinocytes. N=4, unpaired Student's t-test.

**g-h)** Images and quantification of SERPINB5 immunostainings in sgNT1 and sgDPM1 HaCaT keratinocytes. Scale bar indicates 10μm distance. Panel represents 3 biological replicates. Unpaired Student's t-test (N=3). **i)** Western blot showing expression of the resepctive SERPIN-GFP constructs in sgDPM1 HaCaT keratinocytes. GFP used as control. GAPDH used as loading control. Panel shows representative of 3 biological replicates. **j)** Dispase-based dissociation assays to semi quantitatively assess cell-cell adhesion in primary human keratinocytes overexpressing SERPINB5-GFP in sgDPM1 background (N=4), One-way ANOVA, Dunnett's multiple comparision test used for statistics.

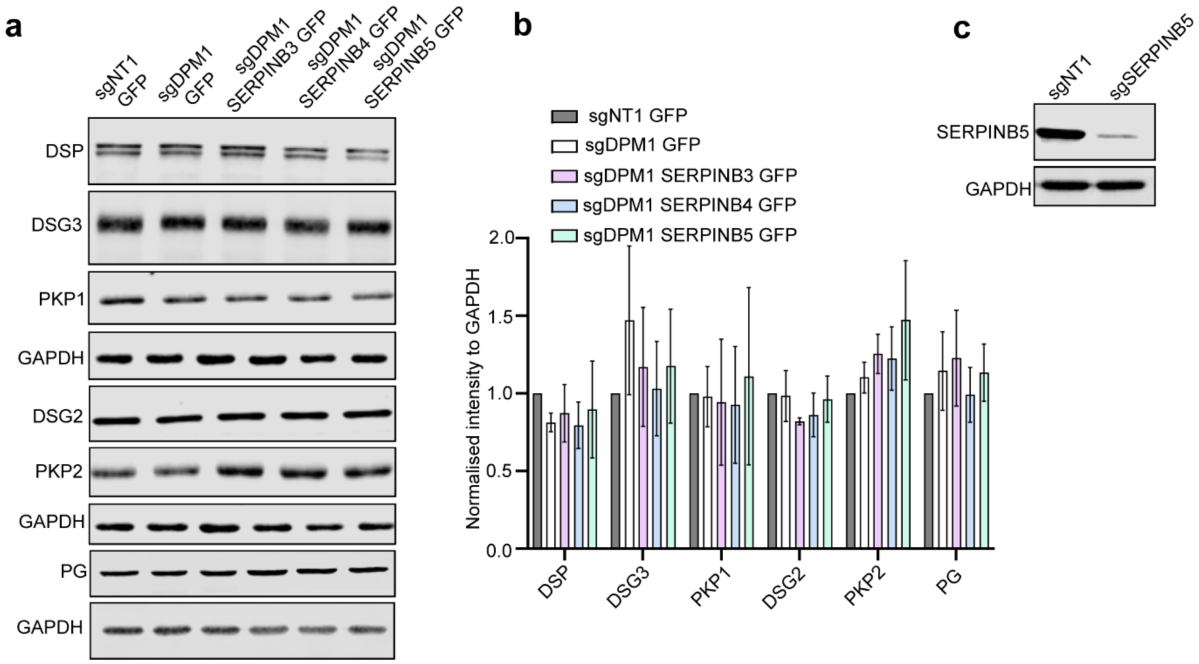


**Figure S3: a-b)** Western blot images and quantifications of desmosomal proteins from sgNT1 and sgDPM1 HaCaT keratinocytes expressing the indicated SERPIN-GFP constructs. GAPDH used as internal loading control (N=3). **c)** Western blot showing SERPINB5 levels in sgNT1 and sgSERPINB5 HaCaT keratinocytes. Image represents 3 biological replicates.

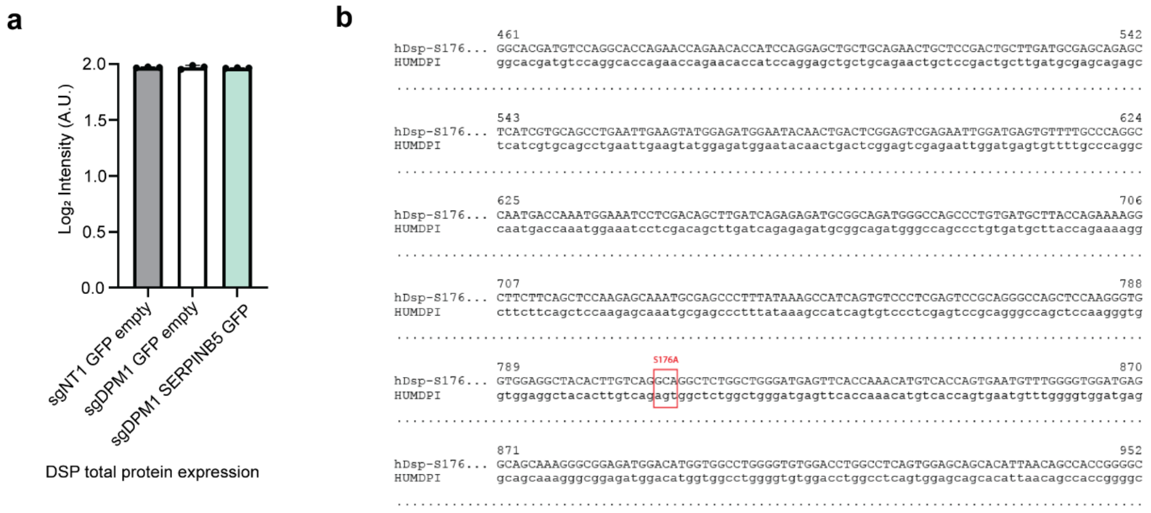


**Figure S4: a)** Graph showing endogenous total DSP protein quantification from mass spectrometry data in sgNT1, sgDPM1 and sgDPM1 SERPINB5 overexpressing cell lines. Y-axis denoted Log2 fold change (FC) of intensity values. Dots represent individual biological replicates. **b)** Sequencing data showing point mutation of Serine at 176 position to Alanine of Dsp (HUMDP1=hDsp1 WT, as template).

**Supplementary table 1: List of primers**

| | |
|---|---|
| sgRNA_hDPM1_23063_for | CACCGTGGAAGCCCAGATGGAACAA |
| sgRNA_hDPM1_23063_rev | AAACTTGTTCCATCTGGGCTTCCAC |
| sgRNA_hDPM2_23063_for | CACCGACATACCAAGAGAATCACCC |
| sgRNA_hDPM2_23063_rev | AAACGGGTGATTCTCTTGGTATGTC |
| sgRNA_hDPM3_23063_for | CACCGAGGACTTCCTGGCAGGACAA |
| sgRNA_hDPM3_23063_rev | AAACTTGTCCTGCCAGGAAGTCCTC |
| sgRNA_hDSG2_5092_for | CACCGCAACGAACCAGTGTTCACAC |
| sgRNA_hDSG2_5092_rev | AAACGTGTGAACACTGGTTCGTTGC |
| sgRNA_SERPINB5_13899_for | CACCGTACGAAGAGACCGTATGCAA |
| sgRNA_SERPINB5 _13899_rev | AAACTTGCATACGGTCTCTTCGTAC |
| sgRNA_DSP_5101_for | CACCGACAACAACGAGCGCAGCAAG |
| sgRNA_DSP_5101_rev | AAACCTTGCTGCGCTCGTTGTTGTC |
| Asc1_hSERPINB3_for | CGCCGCGGCGCGCCATGAATTCACTCAGTGAAGCCAAC |
| hSERPINB3_Xho1_rev | TACAGCCTCGAGCGGGGATGAGAATCTGCCA |
| Asc1_hSERPINB5_for | CGCCGCGGCGCGCCATGGATGCCCTGCAACTAGC |
| hSERPINB5_Xho1_rev | TACAGCCTCGAGAGGAGAACAGAATTTGCCAAAGAAA |
| hDSP S176A_for | CACTTGTCAGGCAGGCTCTGGCTGG |
| hDSP S176A_rev | TAGCCTCCACCACCCTTG |
